# *In cellulo* DNA assembly for targeted genomic integration and rearrangement in human cells

**DOI:** 10.1101/2025.06.16.659926

**Authors:** Sébastien Levesque, Nozomu Kawashima, GueHo Hwang, Basheer Becerra, Vivien A.C. Schoonenberg, William Mannherz, Luke Homfeldt, Luca Pinello, Suneet Agarwal, Daniel E. Bauer

## Abstract

Although therapeutic genome editing holds great potential to remedy diverse inherited and acquired disorders, targeted installation of medium to large sized genomic modifications in therapeutically relevant cells remains challenging. We have developed an approach that permits DNA sequence assembly and integration in human cells leveraging CRISPR-targeted dual flap synthesis. This method, named prime assembly, allows for RNA-programmable site-specific integration of single- or double-stranded DNA fragments. Unlike homology-directed repair, prime assembly was similarly active in dividing and non-dividing cells. We applied prime assembly to perform targeted exon recoding, transgene integration, and megabase-scale rearrangements, including at therapeutically relevant loci in primary human cells. Prime assembly expands the capabilities of genome engineering by enabling the targeted integration of medium to large sized DNA sequences without relying on double-stranded DNA donors, nuclease-driven double strand breaks, or cell cycle progression.

## Introduction

The CRISPR-Cas toolkit provides unprecedented opportunities to engineer the human genome with high precision, efficiency, and versatility. Despite substantial progress, developing safe and effective personalized genome editing therapies requires significant resources, creating obstacles for clinical translation. Since most genetic disorders are caused by heterogeneous mutations, generalizable approaches to replace larger DNA sequences^1^ could provide universal gene-specific solutions to remedy most pathogenic mutations causing a disease. Also, site-specific integration of synthetic transgenes could have numerous applications, such as installing antigen receptors to redirect immune cells for the treatment of malignancies and autoimmune diseases^2–5^.

Therapeutic nuclease-based editing of hematopoietic stem and progenitor cells (HSPCs) has enabled the first CRISPR-Cas9 therapy approved by regulatory agencies in North America and Europe, exagamglogene autotemcel^6–10^. As the field advances towards new therapeutic indications and delivery paradigms, cell context-specific barriers are emerging. The development of gene correction or knock-in strategies based on homology-directed repair (HDR), which occurs in the S and G2 phases of the cell cycle^11,12^, are restricted to actively dividing cells. Nuclease-driven DNA double-strand breaks (DSBs) also generate safety concerns regarding off-target editing^13–16^ and complex genomic rearrangements^17–20^ which are exacerbated in proliferating cells^21,22^. Gene editing technologies that do not require nuclease-driven DSBs and cell cycle progression could overcome these hurdles.

A variety of genome editors have been developed to precisely substitute DNA sequences without requiring DNA DSBs, including base and prime editors^23–25^. Among these technologies, prime editing and click editing utilize CRISPR-targeted flap synthesis to install precise base substitutions and small insertions and deletions^25–27^. Although prime editing compares favorably with other CRISPR tools in terms of product purity, off-target editing, and genotoxicity^25,28–32^, it is currently limited to short- and medium-sized genomic modifications of fewer than 250 base pairs (bp)^33^. A two-step genome editing process, where a prime editor installs a landing pad followed by recombinase-driven DNA integration, can expand this targeting range to large-sized modifications of more than 250 bp, although dependent on complex multicomponent delivery^34–36^. Other recombinase systems depend on systematic discoveries^37^ or protein engineering to achieve target specificity^38^. While promising, reported recombinase- and transposase-based technologies^34–42^ currently rely on double-stranded DNA (dsDNA) or adeno-associated virus (AAV) donors that induce toxicity in hematopoietic cells due to cGAS-STING sensing^43,44^ and p53 activation^45–47^, respectively. Given their lower immunogenicity, single-stranded DNA (ssDNA) and circular ssDNA (cssDNA) donors are better tolerated in many cell contexts^43,48–50^ and could potentially facilitate effective site-specific exon recoding or therapeutic transgene integration.

A new paradigm in molecular cloning emerged in 2009 when Gibson *et al*. reported the enzymatic assembly of kilobase DNA molecules^51^. While this *in vitro* technique quickly became a standard, molecular cloning through *in cellulo* DNA assembly in commonly used *E. coli* strains had been reported more than a decade before, although never gaining popularity^52–54^. DNA assembly in widely used cloning strains (e.g. *E. coli* DH5α) depends on an incompletely characterized RecA-independent recombination (RAIR) mechanism^54–57^. These DNA assembly modalities require common steps: the generation of complementary ssDNA overlaps, homology-directed annealing, fill-in synthesis, and ligation. Whereas RAIR occurs via 5’ ssDNA homology-directed annealing^54^, *in vitro* isothermal Gibson assembly occurs through 3’ ssDNA overlaps^51^. Interestingly, strategies based on paired prime editing guide RNAs (pegRNAs) require similar DNA repair steps^33,34,58–60^, suggesting that human cells may possess the capabilities to permanently incorporate exogenous DNA sequences to their genome via *in cellulo* DNA assembly.

Inspired by RAIR and isothermal Gibson assembly cloning^51–54^, we developed an approach for site-specific DNA assembly and integration in human cells using CRISPR-targeted 3’ flap synthesis. We applied this method, which we term *prime assembly*, to perform targeted exon recoding, transgene integration, and megabase-scale rearrangements at multiple loci using either 3’-overhang dsDNA or ssDNA donors. Using a CDK4/6 inhibitor and cell cycle analysis, we observed efficient prime assembly in cells arrested in G1. We found that prime assembly was active in human primary hematopoietic stem and progenitor cells. Our study establishes a new modality to introduce medium to large genetic modifications to the human genome.

## Results

### Targeted exon recoding using prime assembly

We hypothesized that synthesizing 3’ flaps via prime editing could enable site-specific *in cellulo* DNA assembly and integration of exogenous ssDNA donors, or a dsDNA donor with 3’ overhangs (**Fig. 1a**). After homology-directed donor(s) annealing to the 3’ flaps, endogenous cellular DNA repair pathways could ensure the excision of the unedited duplex, free ssDNA fill-in synthesis, and ligation (**Fig. 1a**). Reminiscent of RAIR and isothermal Gibson assembly cloning^51–54^, we term this approach prime assembly (PA).

**Figure 1.**
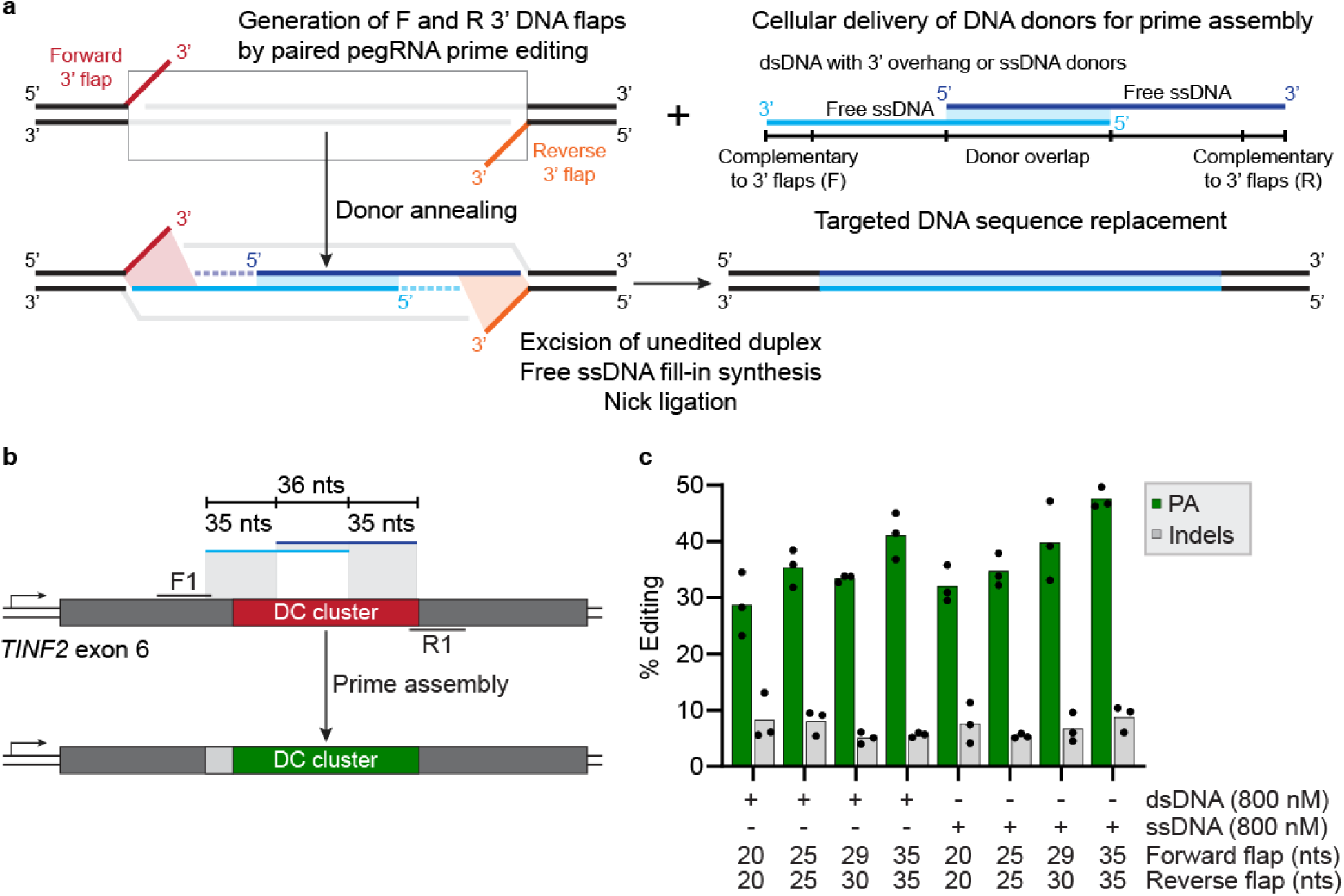
Prime assembly enables sequence replacement in human cells. **(a)** Schematic representation of prime assembly (PA). (**b**) Schematic of *TINF2* DC cluster recoding using PA. (**c**) PA and indel quantification at *TINF2* as determined by CRISPResso2 analysis from amplicon sequencing. K562 cells were electroporated with PA vectors and the indicated concentration of donors with (dsDNA_v5) or without (ssDNA_v5) prior annealing, and genomic DNA was harvested 3 days post-nucleofection. *n* = 3 independent biological replicates. PA defined as perfect targeted integration and indels as all other edits.

As a proof of concept, we first designed a medium-sized therapeutic exon recoding strategy. *TINF2* encodes for the central component of the shelterin telomere protection complex and is the second most frequently mutated gene associated with the telomere biology disorder dyskeratosis congenita (DC)^61,62^. Dominant gain-of-function mutations in a small region of *TINF2* exon 6, which encodes for 30 amino acids known as the DC cluster (**Fig. 1b** and **Supplementary Fig. 1**), result in very short telomeres and more severe symptoms than other DC-causing mutations^63–66^. We reasoned that a single approach could potentially remedy nearly all known pathogenic DC cluster mutations. To prevent DNA flaps and donors annealing to the endogenous genomic sequence, we performed codon optimization throughout the targeted 106-bp region to recode the exon without altering the amino acid sequence (**Fig. 1b** and **Supplementary Fig. 1**). We electroporated K562 cells with plasmids encoding for PEmax^67^ and a pair of epegRNA targeting *TINF2*, along with various concentrations of synthetic ssDNA donors, either annealed (dsDNA) or not (ssDNA) prior to electroporation. We observed up to 42% editing with ssDNA donors, as determined by Sanger sequencing (**Supplementary Fig. 1b**). Electroporating 800 nM of synthetic ssDNA donors protected with three phosphorothioate bonds on both 5’ and 3’ ends, without prior *in vitro* annealing, yielded the highest efficiency (**Supplementary Fig. 1c**).

Our initial design (v1) relied on 25-nucleotide (nts) flaps and no fill-in synthesis. We then tested whether shorter flaps could support PA and observed an abrogation of editing with flaps shorter than 14 nts (**Supplementary Fig. 1 d-f**). We designed ssDNA donors with different complementary overlap lengths (v1-v4) between the two ssDNA donors, and observed similar efficiencies with overlaps of 20 to 56 nts, suggesting that short fill-in synthesis is not a limiting factor (**Supplementary Fig. 1 d-f**), as previously reported for twin prime editing^33,34^. We switched to the PEmax-La (PE7) editor^68^ with standard (La-accessible) pegRNAs, and tested flap lengths ranging from 20 to 35 nts and ssDNA donors sharing 36 nts of overlaps (v5) (**Fig. 1b**). We achieved an average of 47.6% precise editing with 35-nts flaps, as determined by amplicon sequencing (**Fig. 1c**). Characterization of the smaller fraction of unintended indel outcomes revealed the presence of incomplete PA alleles, composed of unidirectional or bidirectional flap integration accompanied by deletion of the sequence located between the flaps (**Supplementary Fig. 2**). PA requires two single-strand breaks (SSBs) and may generate staggered DSB intermediates, leading to unintended insertions and deletions at the target site, as previously reported for prime editing with two nicks^32,67,69,70^. Taken together, PA enables efficient medium-sized exon recoding.

### Prime assembly enables site-specific transgene integration

Efficient HDR has been reported using 3’-overhang dsDNA donors^71^. We reasoned that we could repurpose this type of donor to achieve targeted transgene integration by generating 3’ overhangs that are compatible with 3’ PA flaps. We generated 3’-overhang dsDNA donors by PCR amplification and lambda exonuclease digestion under the protection of phosphorothioate bonds to generate complementary 3’ overhangs (illustrated in **Supplementary Fig. 3**)^71^. We electroporated K562 cells with dsDNA donors and pairs of pegRNAs targeting the *TRAC* locus to generate 32-nts (v1) or 25-nts (v2) flaps. Our leading pair of pegRNAs (F1_v1 + R2_v1) achieved an average PA allele frequency of 4.9%, as determined by ddPCR (**Fig. 2a,b**). We amplified the integration junctions and confirmed precise site-specific integration by Sanger sequencing across all biological replicates (**Supplementary Fig. 4**). We then targeted the EGFP reporter cassette to transcriptionally active loci to confirm functional integration. Under transcriptional control of the endogenous promoter, we observed an average of 7.3% and 8.5% EGFP^+^ cells by targeting the *IL2RG* and *AAVS1* loci, respectively (**Fig. 2c,d** and **Supplementary Fig 5a,b**). We then tested whether we could integrate larger transgene cassettes by incorporating an exogenous human *PGK1 (hPGK1)* promoter and a CD19-CAR to our 3’-overhang dsDNA donor (**Fig. 2e**). After electroporation, we analyzed the percentage of EGFP^+^ cells 21 and 35 days post-nucleofection to eliminate fluorescence signal from non-integrated donors. We observed a higher fluorescence signal with the *hPGK1* promoter (**Supplementary Fig. 5**), and an average of 22.2% and 6.3% EGFP^+^ cells 35 days after nucleofection with the 1.6 kb and 3.1 kb donors, respectively (**Fig. 2 e,f**). Prime assembly supports the integration of larger transgene cassettes, albeit with decreased efficiency, as previously observed for HDR using linear dsDNA donors^72^.

**Figure 2.**
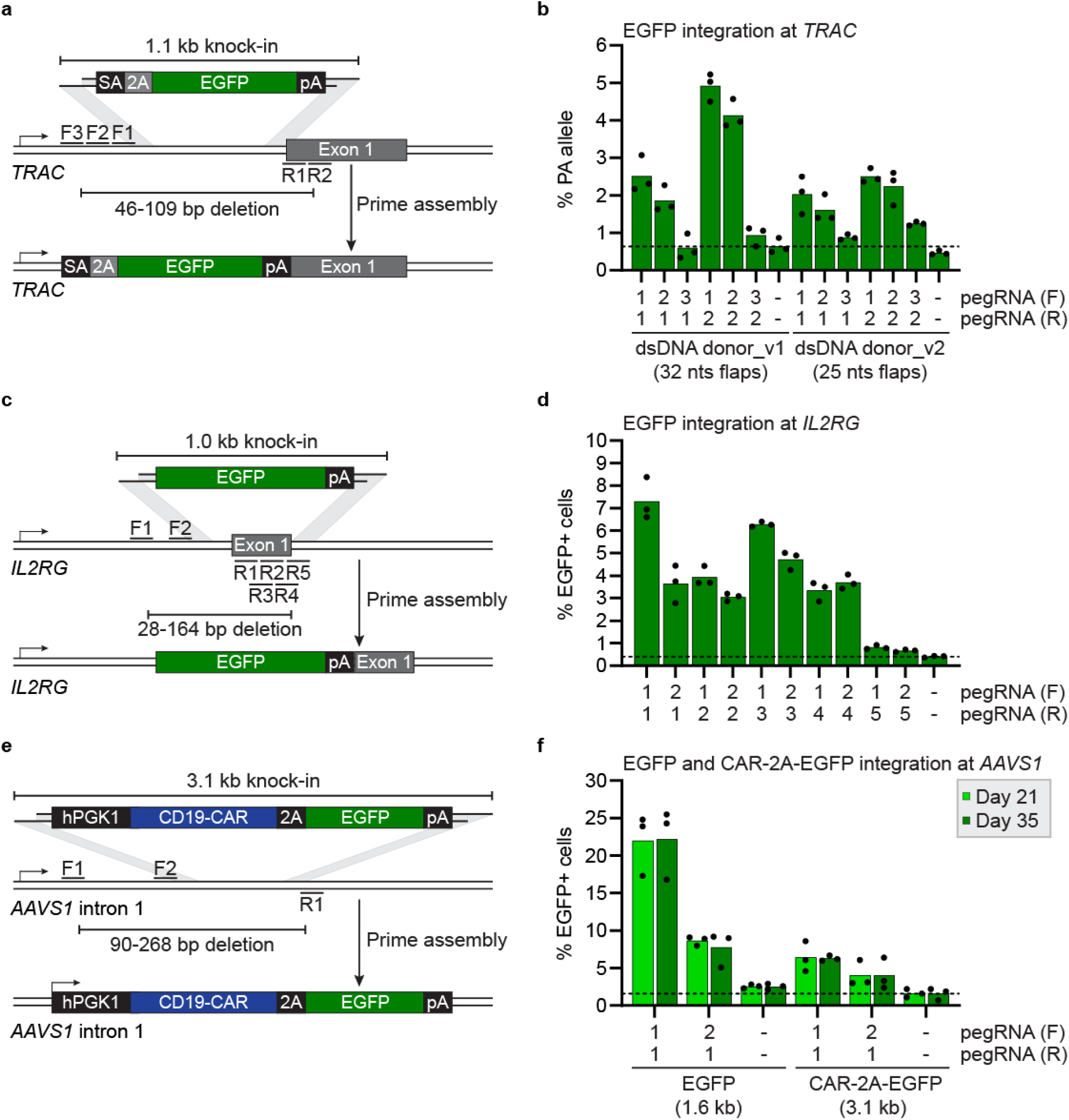
Site-specific transgene integration using 3’-overhang double-stranded DNA donor. (**a**) Schematic representation of targeted transgene integration at the *TRAC* locus using PA. (**b**) Percentage of PA allele as determined by ddPCR. K562 cells were electroporated with PA vectors and 1.5 µg 3’-overhang dsDNA donor targeting *TRAC*, and genomic DNA was harvested three days post-nucleofection. *n* = 3 independent biological replicates. Dotted lines indicate the average background level of the donor only controls. (**c**) Same as in (**a**) for targeted transgene integration at the *IL2RG* locus. (**d**) Percentage of EGFP^+^ cells as determined by flow cytometry. K562 cells were electroporated with PA vectors and 1.5 µg 3’-overhang dsDNA donor targeting *IL2RG*, and the percentage of EGFP^+^ cells was quantified 7 days post-nucleofection. *n* = 3 independent biological replicates. Dotted lines indicate the average background level of the donor only controls (**e**) Same as in (**a**) for targeted transgene integration at the *AAVS1* locus using an equimolar concentration of donor for CAR-2A-EGFP (3.1 kb). (**f**) Percentage of EGFP^+^ cells as determined by flow cytometry. K562 cells were electroporated with PA vectors and 3’-overhang dsDNA donors targeting *AAVS1*, and the percentage of EGFP^+^ cells was quantified 21 and 35 days post-nucleofection. *n* = 3 independent biological replicates. Dotted lines indicate the average background level of the donor only controls

We then tested whether long ssDNA donors could support site-specific transgene integration at *AAVS1* and *IL2RG*. We electroporated K562 cells with long ssDNA donors sharing overlapping regions of 500 bp (v1) or 100 bp (v2) and observed an average of 2.8% and 3.3% EGFP^+^ cells, respectively (**Fig. 3a,b**). In contrast, an average of 10.5% EGFP^+^ cells was observed when electroporating an equimolar ratio of 3’-dsDNA donor, a 3.2-fold increase over ssDNA donors (v2) (**Fig. 3b**). Increasing ssDNA molarity modestly increased integration efficiencies at *AAVS1* and *IL2RG*, and efficiencies remained lower than 3’-overhang dsDNA donor (**Supplementary Fig. 6**). We then used an asymmetric design with a long ssDNA donor and a short synthetic ssDNA sharing an overlapping region of 50 bp (v3) (**Fig. 3c**). Using this design at *IL2RG*, we achieved an average of 3.5% EGFP^+^ cells with ssDNA, a 1.6-fold decrease over the corresponding 3’-overhang dsDNA donor (**Fig. 3d**). The capacity of PA to accommodate long ssDNA donors with short overlapping regions opens opportunities for targeted therapeutic transgene integration in primary hematopoietic cells that are sensitive to dsDNA^43,44^.

**Figure 3.**
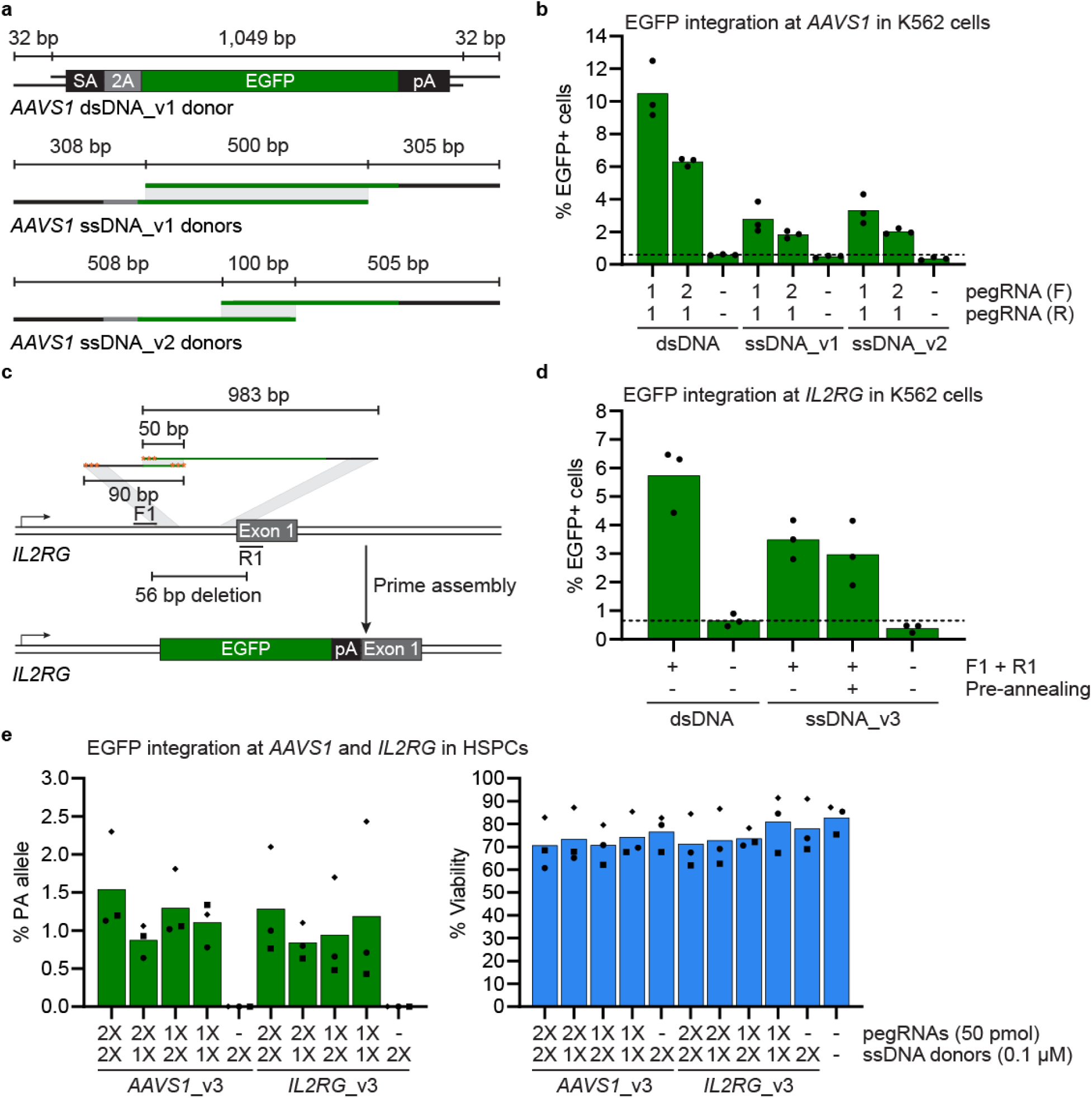
Site-specific transgene integration using single-stranded DNA donors in K562 cells and CD34^+^ HSPCs. (**a**) Schematic representation of targeted transgene integration at the *AAVS1* locus using two different ssDNA donor designs and pegRNAs encoding for 32-nts flaps. (**b**) Percentage of EGFP^+^ cells as determined by flow cytometry. K562 cells were electroporated with PA vectors and 200 nM ssDNA donors targeting *AAVS1* or an equimolar concentration of 3’-overhang dsDNA donor, and the percentage of EGFP^+^ cells was quantified 7 days post-nucleofection. *n* = 3 independent biological replicates. Dotted lines indicate the average background level of the donor only controls. (**c**) Same as in (**a**) for site-specific integration at *IL2RG*. Stabilizing phosphorothioate bonds are illustrated with orange stars. (**d**) Percentage of EGFP^+^ cells as determined by flow cytometry. K562 cells were electroporated with PA vectors and 200 nM ssDNA donors targeting *IL2RG* or an equimolar concentration of 3’-overhang dsDNA donor, and the percentage of EGFP^+^ cells was quantified 7 days post-nucleofection. *n* = 3 independent biological replicates. Dotted lines indicate the average background level of the donor only controls. (**e**) Percentage of PA allele as determined by ddPCR, and cell viability as determined by trypan blue staining and manual cell counting. A total of 2.5x10^5^ CD34^+^ HSPCs were electroporated with PE mRNA, and the indicated concentration of synthetic pegRNAs and ssDNA donors. Genomic DNA was collected three days post-nucleofection. *n* = 3 independent biological replicates performed with HSPCs from three different donors. Circle, donor 1; Diamond, donor 2; Rectangle, donor 3.

We have previously achieved highly efficient prime and twin prime editing in CD34^+^ human hematopoietic stem and progenitor cells (HSPCs) by modulating nucleotide metabolism^73^. Using our optimized protocol^73^, we electroporated CD34^+^ HSPCs from three different healthy donors with PE7 mRNA and various concentrations of pegRNAs and ssDNA donors to target an EGFP reporter to the *AAVS1* and *IL2RG* loci. With 100 pmol pegRNAs and 200 nM ssDNA donors, we observed 60.8%-82.9% viability 24 hours post-nucleofection, and an average of 1.5% and 1.3% PA allele at the *AAVS1* and *IL2RG* loci, respectively (**Fig. 3e**). These results demonstrate that *in cellulo* DNA assembly is applicable in therapeutically-relevant hematopoietic cells.

### Multi-fragment assembly and megabase deletions

A key benefit of DNA assembly cloning methods^51–54^ is their capacity to assemble multiple fragments together. We reasoned that PA could also assemble more than two ssDNA fragments in human cells under selective pressure. To test this, we designed an intron nesting approach at the *ATP1A1* locus^70,74^ to assemble four ssDNA fragments encoding for a U6-pegRNA cassette while installing a dominant gain-of-function mutation (*ATP1A1*-T804N^70^) conferring resistance to ouabain, a plant-derived inhibitor of the sodium-potassium pump (**Supplementary Fig. 7**). We electroporated K562 cells with PA vectors and four ssDNA fragments, and treated cells with 0.5 µM ouabain three days post-nucleofection until all non-resistant cells were eliminated. We performed PCR-based genotyping and detected the expected PA allele in the bulk population of resistant cells (**SupplementaryFig. 7c**). To confirm the functional integration of the *B2M*-L7Stop_v3^73^ pegRNA reporter, we re-electroporated K562 cells with a PE7-expressing plasmid and observed an average of 27% edited allele at the *B2M* locus three days post-nucleofection, as determined by Sanger sequencing (**Supplementary Fig. 7d**). We readily cloned the PA allele into a plasmid, confirmed precise assembly and integration of the four fragments via Sanger sequencing, and used the vector to generate a long ssDNA donor in-house. Using two ssDNA donors, we achieved higher integration efficiencies and observed an average of 61% editing at the reporter *B2M* locus (**Supplementary Fig. 7e,f**). Our results confirmed that prime assembly can assemble and integrate up to four ssDNA fragments in human cells.

Paired pegRNA approaches, such as twin prime editing and PRIME-Del, enable deletions of less than 10 kb^33,34,60^. We reasoned that PA could excise larger genomic sequences while installing a selection marker. We designed pairs of pegRNAs to integrate a puromycin selection marker while introducing deletions of 268 bp, 10 kb, and 1 Mb on chromosome 19 (**Fig. 4a**). Three days after nucleofection, we treated cells with 1 µg/ml puromycin until all non-resistant cells were eliminated. We observed a marked increase in the percentage of PA alleles after puromycin selection, as determined by ddPCR (**Fig. 4b**). As expected, targeting efficiencies were inversely proportional to the distance between the cut sites (**Fig. 4b**). We performed long-range PCR-based genotyping with primers binding outside the PA junctions and readily detected the expected 1 Mb deletion allele (**Fig. 4d**). We also performed long-read nanopore sequencing and confirmed the expected 10 kb and 1 Mb PA alleles across independent biological replicates. We then designed additional pairs of pegRNAs to install kilobase and megabase deletions on chromosome 7 (**Supplementary Fig. 8a**). We achieved large deletions on chromosome 7 without pegRNA optimization, as confirmed by ddPCR, PCR-based genotyping, and long-read nanopore sequencing across independent biological replicates (**Supplementary Fig. 8**).

**Figure 4.**
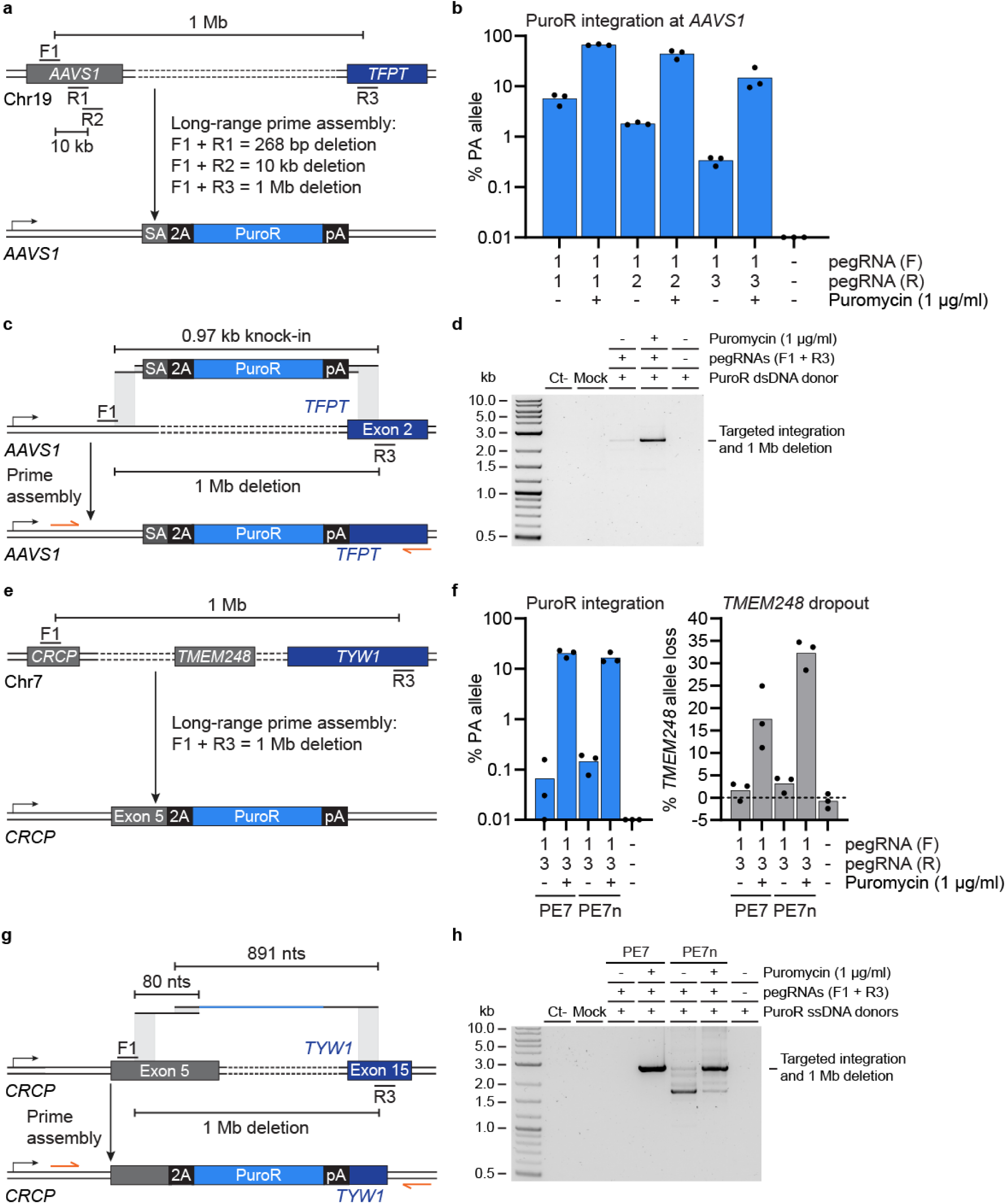
Long-range prime assembly enables megabase deletion in human cells. (**a**) Schematic representation of targeted deletions and the integration of a puromycin-resistance (PuroR) cassette at chromosome 19 via long-range prime assembly. (**b**) Percentage of PA allele as determined by ddPCR. K562 cells were electroporated with PA vectors and 1.5 µg 3’-overhang dsDNA donor targeting *AAVS1*, and cells were cultured with 1 µg/ml puromycin three days post-nucleofection until all non-resistant cells were eliminated. *n* = 3 independent biological replicates. (**c**) Schematic representation of targeted PuroR transgene integration and one megabase deletion at chromosome 19. (**d**) PCR-based genotyping of the expected PuroR transgene integration and megabase deletion allele after prime assembly and puromycin selection, as described in (**b**). Representative gel is from one out of three independent biological replicates. (**e**) Same as in (**a**) for targeted megabase deletion and PuroR integration at chromosome 7. (**f**) Percentage of PA allele and *TMEM248* copy loss as determined by ddPCR. K562 cells were electroporated with PA or nuclease PA vectors and 200 nM ssDNA donors targeting chromosome 7, and cells were cultured with 1 µg/ml puromycin three days post-nucleofection until all non-resistant cells were eliminated. *n* = 3 independent biological replicates. (**g**) Same as in (**c**) for targeted megabase deletion at chromosome 7 using ssDNA donors. (**h**) Same as in (**d**) for targeted PuroR integration and megabase deletion at chromosome 7. Representative gel is from one out of three independent biological replicates.

Previous studies have reported low frequencies of off-target dsDNA donor integrations^71,72^. In line with these results, we observed puromycin-resistant cells in our donor only controls. We reasoned that ssDNA donors could decrease these off-target integration events and facilitate the enrichment of cells with the intended megabase-scale deletions. We also hypothesized that nuclease-driven prime assembly, in a process reminiscent of PEDAR^59^, could facilitate large-scale chromosomal rearrangements. We electroporated cells with PE7 or PE7 nuclease (PE7n) vectors and two ssDNA donors, and successfully introduced a megabase-scale deletion on chromosome 7 with little to no puromycin-resistant cells in our donor only controls (**Fig. 4e-h**). Importantly, we observed an average of 17.5% and 32.2% *TMEM248* allele loss, a gene present in the intended 1 Mb deletion, in bulk populations of resistant cells after PA and nuclease PA, respectively (**Fig. 4f**). We readily detected the 1 Mb deletion allele by long-range PCR and confirmed the expected allele by long-read nanopore sequencing (**Fig. 4h**). We then tested whether we could achieve targeted chromosome arm deletion by installing a 93 Mb deletion on chromosome 7 (**Supplementary Fig. 9a**). While no chromosome 7 q arm deletion was observed with standard PA, nuclease-driven PA enabled the installation of the 93 Mb deletion, as confirmed by ddPCR, PCR-based genotyping, and long-read nanopore sequencing (**Supplementary Fig. 9a-d**). Finally, we achieved targeted 1 Mb inversion at chromosome 7 using nuclease PA (**Supplementary Fig. 9 e,f**). These results suggest that nuclease-induced DSBs facilitate chromosome-scale deletions and megabase inversion, although at the cost of lower product purity. Taken together, PA offers a new tool for installing large-scale genomic rearrangements for functional genomics applications in human cells.

### Prime assembly occurs in G1-arrested cells

The dependence of HDR on cell cycle progression limits its applications in primary cells, many of which are quiescent or post-mitotic. We designed an assay to assess whether prime assembly could occur in cells arrested in G1 using the CDK4/6 inhibitor palbociclib (PD 0332991, hereafter referred to as PD). We treated K562 cells for 24 hours with 5 µM PD, the lowest dose that enabled G1 arrest in ≥ 90% of cells with no impact on viability, and observed a marked decrease in cells progressing through S and G2/M, as determined by Hoechst DNA staining and flow cytometry analysis (**Fig. 5 a-c**). We note that a fraction of K562 cells can progress through S and G2/M after more than 24 hours of PD treatment (**Fig. 5c**), possibly due to fractional resistance^75^. Treatment with 5 µM PD 24 hours before nucleofection and/or 72 hours after nucleofection abrogated HDR at *AAVS1* from an average of 17% to below the level of detection by Sanger sequencing (**Supplementary Fig. 10**). We then treated K562 cells for 24 hours with 5 µM PD, electroporated cells with plasmid encoding PE7 or PE7n, pegRNAs, and a 3’-dsDNA overhang donor to integrate an EGFP reporter at the *AAVS1* and *IL2RG* loci. After nucleofection, we enabled cell cycle progression (without PD) or kept cells arrested in G1 (5 µM PD) for 72 hours before genotyping (**Fig. 5a**). While untreated cells expanded 3.1-to 4.8-fold after three days of culture, minimal expansion was observed with PD-treated cells (**Fig. 5d**). We quantified the percentage of PA allele by ddPCR in untreated and PD-treated cells and observed little to no impact on EGFP integration using either PE7 or PE7n (**Fig. 5d**). These results suggest that prime assembly can occur in G1-arrested cells with or without nuclease-driven DSBs.

**Figure 5.**
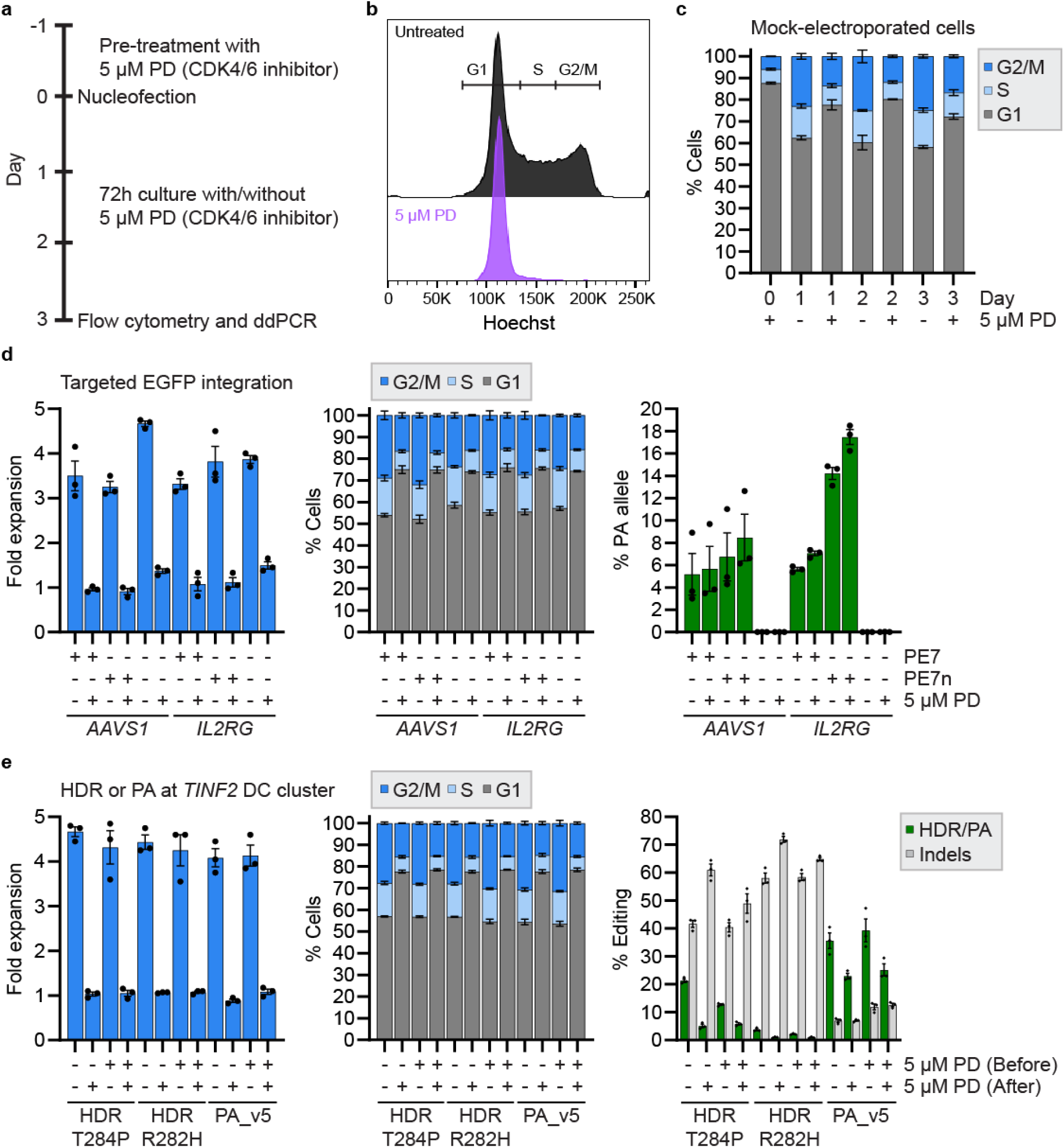
Prime assembly occurs in G1-arrested cells. (**a**) Timeline for the cell cycle experiment. K562 cells are pre-treated for 24 hours with 5 µM PD 0332991, electroporated, and cultured for three days in the presence or absence of 5 µM PD 0332991. (**b**) Representative flow cytometry plot for cell cycle analysis. K562 cells were cultured for 24 hours in the presence or absence of 5 µM PD 0332991, and cell cycle progression was measured by Hoechst DNA staining and flow cytometry. Representative image is from one of four independent biological replicates. (**c**) Cell cycle progression as determined by flow cytometry. K562 cells treated for 24 hours with 5 µM PD 0332991 were mock electroporated and cultured in the presence or absence of 5 µM PD 0332991. Cell cycle progression was monitored for four days. Data are plotted as mean ± s.e.m. from *n* = 3 independent biological replicates. (**d**) Same as in (**c**) with K562 cells electroporated with PA vectors and 3’-overhang dsDNA donors targeting *AAVS1* or *IL2RG* for EGFP transgene integration. Cell cycle analyses, measurement of fold expansion, and ddPCR-based genotyping were performed three days post-nucleofection. Data are plotted as mean ± s.e.m. from *n* = 3 independent biological replicates. (**e**) Impact of cell cycle arrest on HDR and prime assembly. K562 cells were cultured in the presence or absence of 5 µM PD 0332991 (before electroporation), electroporated with HDR or PA vectors and ssDNA donors targeting *TINF2*, and cultured for three days in the presence or absence of 5 µM PD 0332991 (after electroporation). Cell cycle analysis, measurement of fold expansion, and genomic DNA purification were performed three days post-nucleofection. The percentage of edited allele was determined by amplicon sequencing. Data are plotted as mean ± s.e.m. from *n* = 3 independent biological replicates.

We then installed dyskeratosis congenita mutations via HDR (*TINF2*-T284P or R282H) or recoded the DC cluster via prime assembly at the *TINF2* locus. We observed a marked decrease in HDR efficiency with R282H compared to T284P using the same sgRNA (**Fig. 5e**), corroborating the inverse relationship between mutation incorporation and the distance from the cut site when using an ssDNA donor^76^. Treatment with 5 µM PD before and after electroporation decreased HDR efficiency by 3.7- and 3.9-fold for *TINF2*-T284P and *TINF2*-R282H, respectively (**Fig. 5e**). For prime assembly, we observed a 1.4-fold decrease in editing efficiency in PD-treated cells, and a 1.7-fold increase in indels, suggesting that although PA can occur in G1-arrested cells, some DNA repair factors involved in precise ssDNA assembly and integration might be limiting in G1. Importantly, in actively dividing cells, we observed 10- and 80-fold increases in precise editing-to-indels ratio with PA as compared to HDR for *TINF2*-T284P and R282H, respectively (**Fig. 5e**). In cells treated with PD before and after electroporation, we observed 17- and 134-fold increases in precise editing-to-indels ratio with PA as compared to HDR for *TINF2*-T284P and R282H, respectively (**Fig. 5e**). Together, our results suggest that prime assembly outperforms HDR in non-proliferating cells while facilitating medium-sized recoding using ssDNA donors.

## Discussion

The ability to assemble DNA sequences *in vitro*^51^ emerged as a new paradigm for molecular cloning. By leveraging CRISPR-targeted 3’ flap synthesis and endogenous cellular DNA repair pathways, we demonstrate that this modality can extend to human cells. While molecular cloning via *in vitro* or *in cellulo* DNA assembly relies on exonuclease degradation to generate complementary 3’ or 5’ ssDNA overhangs^51,54^, prime assembly depends on DNA synthesis, a programmable framework for precision genome editing. Importantly, prime assembly can also accommodate ssDNA donors, expanding the versatility of the approach for therapeutic genome editing and functional genomics.

We observed higher editing efficiencies with shorter genomic modifications, which enable delivery of higher concentrations of DNA donors. Our observations suggest that the local concentration of donors available for homology-directed annealing to the 3’ PA flaps may be limiting when delivering larger DNA donors. A promising future direction may be the development of strategies to recruit DNA donors to the PA site to facilitate DNA assembly and integration.

The installation of large-scale structural genomic modifications in human cells provides opportunities to haploidize specific genomic loci or model diseases^77–79^. To the best of our knowledge, this is the first report of megabase-scale deletions achieved in human cells using a prime editor without requiring nuclease-driven DSBs. Coupling prime assembly with a nuclease prime editor also enabled megabase inversion and targeted chromosome arm deletion, expanding the breadth of applications for functional genomics. While we observed benefits with nuclease PA for chromosomal-scale rearrangements, we recommend standard nickase-based PA for targeted genomic integration to maximize product purity.

We successfully programmed targeted integrations of transgenes to two loci in CD34^+^ HSPCs from different healthy donors, demonstrating a novel method to introduce medium- to large-sized modifications in therapeutically relevant hematopoietic cells. Due to their lower immunogenicity^43,44^, ssDNA donors could be used to recode exonic mutation hotspots or target a functional transgene copy to complement a defective gene at its endogenous locus, providing universal strategies to remedy sundry pathogenic mutations. Recoding exons, such as the *TINF2* DC cluster, via HDR would not be possible using short synthetic ssDNA donors due to the long distance between the cut site and the peripheral mutations^76^. Given that *ex vivo* culture with cytokines and cell cycle progression, which are essential for HDR^11,12,80^, correlate negatively with HSC engraftment potential^81,82^ and increase genotoxicity^21,22^, PA may provide an alternative method for medium- to large-sized therapeutic edits. As PA can accommodate synthetic ssDNA donors, unique molecular indexes could also be introduced for long-term clonal tracking^83^. This approach could overcome AAV-induced toxicity^45–47^ as well as imprecise concatemeric AAV vector integration at nuclease DSBs^84,85^. We envision that prime assembly should also enable site-specific antigen receptor transgene integration at endogenous loci for tailored immunity^2–4^. Multiplexed prime assembly coupled with standard prime editing could further improve the functionality of immune cells for cancer immunotherapy (e.g. CAR T cells) by modifying or inactivating key targets^86–88^, without requiring multiple nuclease-induced DSBs. Furthermore, PA could permit targeted integration in post-mitotic cells.

An optimal method for therapeutic targeted integration of gene-sized DNA sequences would avoid the cytotoxicity of dsDNA donors, bypass the genotoxicity and imprecision of nuclease-driven DSBs, and maintain activity in non-dividing cells. To our knowledge, prime assembly is the only currently described method that meets these criteria. In addition, prime assembly only requires a single protein effector. Taken together, prime assembly enables *in cellulo* DNA assembly for precision human genome engineering.

## Methods

### Cell culture and nucleofection

K562 cells were obtained from the ATCC (CCL-243) and cultured at 37°C under 5% CO_2_ in RPMI media supplemented with 10% FBS, and Penicillin/Streptomycin. For each nucleofection, 2x10^5^ cells were electroporated with 750 ng pCMV_PE7^68^ (Addgene 214812), 250 ng of each standard pegRNA^69^ vector (derived from Addgene 132777), and the indicated concentration of prime assembly donor with an Amaxa 4D-nucleofector™ (Lonza) using the SF cell line nucleofection kit (Lonza, catalog no. V4XC-2032) (pulse FF-120). For initial experiments (**Supplementary Fig. 1**), K562 cells were electroporated with 750 ng pCMV PEmax^67^ (Addgene 174820), 250 ng of each tevopreq1-epegRNA^89^ vector (derived from Addgene 174038) harboring the (F+E) scaffold modifications^90^, and the indicated concentration of prime assembly donor. For nuclease prime assembly experiments, the HNH domain of PE7 (Addgene 214812) was restored to generate pCMV_PE7nuclease via Gibson assembly. For HDR experiments, K562 cells were electroporated with 750 ng pX330-U6-Chimeric_BB-CBh-hSpCas9^91^ (Addgene 42230) expressing the sgRNA of interest and 800 nM ssDNA donor. StemSelect PD-0332991 (Sigma, catalog no. 5304870001) was dissolved at 10 mM in water and stored at -80 °C. Ouabain octahydrate (Sigma, catalog no. O3125-250GM) was dissolved at 5 mg/ml in hot water, and working dilutions were prepared in water and stored at -20 °C. Puromycin (Sigma, catalog no. P8833-25MG) was dissolved at 1 mg/ml in water and stored at -20 °C.

Cryopreserved human CD34^+^ HSPCs from mobilized peripheral blood of deidentified healthy donors were obtained from the Fred Hutchinson Cancer Research Center (Seattle, Washington). CD34^+^ HSPCs were cultured in X-Vivo-15 media (Lonza, catalog no. 04-418Q) supplemented with 100 ng/ml human Stem Cell Growth Factor (SCF) (R&D Systems, catalog no. 255-SC-010), 100 ng/ml human thrombopoietin (TPO) (Peprotech, catalog no. 300-18), and 100 ng/ml recombinant human FMS-like Tyrosine Kinase 3 Ligand (Flt3-L) (Peprotech, catalog no. 300-19). CD34^+^ HSPCs were thawed and cultured for 24 hours in the presence of cytokines, and electroporated using the P3 Primary Cell X kit S (Lonza, catalog no. V4XP-3032) according to manufacturer’s recommendations. 2.5 x 10^5^ cells were electroporated with 2000 ng PE7 mRNA^68^, an equimolar ratio of SIV-Vpx mRNA^73^, and the indicated concentration of each pegRNA and ssDNA donors using pulse code DS-130. Following electroporation, 80 µl of media supplemented with cytokines was added to each well and cells were incubated for 10 minutes prior to transfer to the culture plate. Cells were cultured in a 48-well plate in a final volume of 500 µl of media supplemented with 50 µM of each deoxynucleoside^73^. Cell viability was assessed 24 hours post-nucleofection via Trypan Blue staining and manual counting using a hemocytometer, and genomic DNA was purified three days post-nucleofection. Deoxynucleosides (dA, Sigma-Aldrich, catalog #D8668) (dG, Sigma-Aldrich, catalog # D0901) (dC, Sigma-Aldrich, catalog #D0776) (dT, Sigma-Aldrich, catalog #T1895) were resuspended in water at 12.5 mM each, filter-sterilized, and stored at -20°C.

### Prime assembly donors

Short single-stranded DNA (ssDNA) donors were synthesized as ultramers (IDT) at a 4 nmol scale with 5’ phosphorylation. To generate double-stranded DNA donors, ssDNA ultramers were mixed in 50 mM NaCl, 10 mM Tris-HCl (pH 8.0), 1 mM EDTA, and annealed by heating the solution to 95°C for 10 minutes, followed by gradual cooling on a thermocycler. The ssDNA and annealed dsDNA donors were then diluted in IDTE buffer and stored at -20°C. Double-stranded DNA (dsDNA) donors with 3’ overhangs were generated via exonuclease digestion, as previously described^71^. Briefly, donors were amplified from plasmids using Kapa-HiFi polymerase (Roche, catalog no. 07958897001) with primers harboring 5’ phosphorylation, and the expected overhang sequence followed by five consecutive phosphorothioate linkages to block Lambda exonuclease from digesting the donor further. PCR products were purified using SPRIselect beads (Beckman Coulter, catalog no. B23318), digested with Lambda exonuclease (NEB, catalog no. M0262S), and purified again using SPRIselect beads. Donor concentration, purity, and integrity was assessed by nanodrop and agarose gel electrophoresis.

For experiments requiring long ssDNA, donors were amplified using Kapa-HiFi polymerase with a primer harboring a 5’ biotin modification for the DNA strand to separate, and a primer harboring a 5’ phosphorylation for the DNA strand to isolate, as previously described^49^. PCR amplicons were purified with SPRIselect beads. The single strand of interest was then purified via magnetic separation using Streptavidin C1 Dynabeads (Thermo Fisher, catalog no. 11205D). Briefly, Streptavidin C1 Dynabeads were washed two times, mixed with biotinylated PCR amplicons, and incubated at room temperature for 30 minutes with agitation. For magnetic separation, Dynabeads coated with biotinylated amplicons were washed twice, and the supernatant was removed and replaced with 0.125 M NaOH melt solution (prepared fresh) to denature the dsDNA. The solution was placed back on the magnet and the supernatant containing the nonbiotinylated strand was removed gently and mixed immediately with Neutralization buffer (freshly prepared by mixing 100 µl 3M sodium acetate pH 5.2 with 4.8 ml 1X TE buffer). A second round of denaturation and elution was performed with 0.125 M NaOH melt solution using the same neutralization tube. Resulting ssDNA was purified using SPRIselect beads, eluted in IDTE buffer, and ssDNA concentration and purity was assessed by nanodrop. Alternatively, long ssDNA donors were provided by Genscript, resuspended in IDTE buffer, and stored at -20 °C. The EGFP and puromycin resistance transgenes were amplified from *AAVS1*_Puro_hPGK1_EGFP_Donor (Addgene 178088)^70^. Alternatively, the EGFP cassette was cloned in a pUC19 backbone with a splicing acceptor (SA) and a self-cleaving 2A peptide (2A) in-frame with *TRAC* or *AAVS1*. Donor sequences used in this study are provided in the supplementary material, and plasmid vectors will be available at Addgene at the time of publication.

### In vitro transcription and pegRNA synthesis

The PE7 transcription template vector^68^ (Addgene 223022) was linearized using BbsI-HF (NEB, catalog no. R3539L), and mRNA was transcribed using the HiScribe T7 high yield RNA kit (NEB, catalog no. E2050S) using N1-methylpseudouridine (Trilink, catalog no. N-1081) instead of uridine, and co-transcriptional capping with CleanCap AG (Trilink, catalog no. N-7113). The PE7 *in vitro* transcription plasmid template derives from a previously reported vector^73,92^ and encodes for a T7 promoter, a minimal 5’-UTR, a PE7 cassette^68^ harboring a silent mutation disrupting a restriction site for the linearizing BbsI enzyme, a 2x *HBB* 3’ UTR, and a 80-90 bp poly(A) sequence. For SIV Vpx mRNA, the template was generated as previously described^73^. Briefly, the Vpx-SIV vector template^73^ (Addgene 216792) was PCR amplified with a forward primer that correct a T7 promoter inactivating mutation and a reverse primer that appends a 119-nt poly(A) tail to the 3’ UTR. Following IVT, mRNAs were purified using the Monarch RNA Cleanup kit (500 µg) (NEB, catalog no. T2050L) and eluted in 1X nuclease-free IDTE buffer (10 mM Tris, 0.1 mM EDTA, pH 7.5). The mRNA concentration was quantified using Qubit RNA high sensitivity (HS) kit (ThermoFisher, catalog no. Q32852). Synthetic pegRNAs were provided by Integrated DNA Technologies (IDT) and resuspended at 200 pmol/µl in nuclease-free IDTE buffer (10 mM Tris, 0.1 mM EDTA, pH 7.5). The pegRNAs contained 2’-O-methyl modifications and phosphorothioate linkages. All pegRNA sequences and chemical modifications are provided in the supplementary material.

### Genotyping

Genomic DNA was harvested 3 days post-nucleofection using QuickExtract DNA extraction solution (Fisher Scientific, catalog no. NC9904870) following manufacturer’s recommendations. For Sanger sequencing, primers were designed to amplify a 600-800 bp amplicon^93,94^. PCR amplifications were performed with 30 cycles of amplification with Phusion high-fidelity polymerase (NEB, catalog no. M0531L). PCR product quality was evaluated by agarose gel electrophoresis, and purification was performed with SPRIselect beads before Sanger sequencing. Sequencing trace quality was manually expected using Geneious R11 software, and lower quality reactions with background noise were repeated. The percentage of prime assembly alleles and indels were quantified using ICE^95^ (HDR settings) and TIDE^93^ webtools from Sanger sequence data files, respectively. For prime editing at *B2M*, the percentage of prime edited alleles and indels were quantified using BEAT^94^ and TIDE^93^ webtools from Sanger sequence data files, respectively.

For amplicon sequencing, primers were designed to amplify a 200-250 bp amplicon. PCR amplifications were performed with Phusion high-fidelity polymerase, and amplicons were purified with SPRIselect beads. PCR product quality was assessed via agarose gel electrophoresis. Indexing (PCR 2) was performed with 1 µl of locus-specific PCR product. Following beads purification, PCR product quality was assessed by electrophoresis and TapeStation using a DS1000 High Sensitivity ScreenTape assay (Agilent, catalog no. 5067-5585), and quantified with a Qubit dsDNA High Sensitivity (HS) assay kit (Thermo Fisher Scientific, catalog no. Q33231). Amplicons were sequenced using paired-end 150-bp reads in-house on an Illumina MiniSeq system or on an Illumina NovaSeq X system by Novogene (Durham, North Carolina, USA). The percentage of prime assembly, homology-directed repair, or indel alleles was quantified using CRISPResso2^96^. For precise prime assembly or HDR quantification, CRISPResso2 was run in HDR mode using the desired allele as the expected allele. The percentage of indels was determined as the percentage of non-homologous end joins reads plus the percentage of imperfect HDR reads.

For long read nanopore sequencing, primers were designed to amplify a 2-3 kb amplicon encompassing the prime assembly junctions (10 kb and 1 Mb deletions) and the selection marker cassette. PCR amplifications were performed with Phusion high-fidelity polymerase, and amplicons were purified with SPRIselect beads. PCR product quality was assessed via agarose gel electrophoresis. Long read nanopore sequencing was performed by Plasmidsaurus. Uncropped scans of all gels from this study are provided in **Supplementary Fig. 11**. Primers used in this study are provided in the supplementary material.

### Droplet digital PCR

Genomic DNA was extracted and purified using the Monarch spin gDNA extraction kit (NEB, catalog no. T3010L). For each ddPCR reaction, 50 ng of genomic DNA was used, and all conditions were performed in technical triplicates. The droplets were generated using a Bio-Rad QX200 AutoDG droplet digital PCR system with ddPCR supermix (no dUTP) (Bio-Rad, catalog no. 186-3025), and HindIII-HF was supplemented (NEB, catalog no. R3104L) in each reaction. Following droplet generation, samples were amplified using the following conditions: 95°C for 10 min, 40 cycles of 94 °C for 30 s, annealing (56-59 °C) for 60 s, and a final incubation at 98 °C for 10 min. Samples were then kept at 4 °C until analysis. Results were analyzed using the QuantaSoft software, and the percentage of prime assembly (PA) alleles harboring the targeted transgene integration was determined as the ratio of PA allele relative to a genomic reference. Primers and probes used during this study are available in the supplementary material.

### Flow Cytometry and cell cycle analysis

The percentage of EGFP^+^ fluorescent cells was quantified using a BD LSRII flow cytometer, and 1 x 10^5^ cells were analyzed for each condition. Cells were cultured for 7 days post-nucleofection, and donor only conditions were used as a negative control. For experiments using the hPGK1 promoter, cells were cultured for 21 to 35 days to eliminate background fluorescence signal from non-integrated donor. For cell cycle analysis, cells were cultured in the presence or absence of 5 µM PD-0332991 24 hours before and 72 hours after nucleofection. For each nucleofection, 1x10^6^ K562 cells were electroporated with an Amaxa 4D-nucleofector™ (Lonza) using the SF cell line nucleofection kit (pulse FF-120). The fold expansion was measured three days post-nucleofection by Trypan blue staining and manual counting using a hemocytometer. Cells were washed once with PBS, resuspended at 1 x 10^6^ cells/ml in PBS supplemented with 10 µg/ml Hoechst 33342 (Sigma, catalog no. B2261), and stained for 45 minutes at 37 °C in the dark, mixing every 15 minutes. After staining, cells were washed and resuspended in PBS, and 1 x 10^5^ cells were analyzed for each condition using a BD LSRII flow cytometer. Flow cytometric data visualization and analysis was performed using FlowJo (v10).

## Data Availability

All amplicon sequencing data generated during this study will be publicly accessible from the National Center for Biotechnology Information BioProject database at the time of publication.

## Acknowledgements

D.E.B. was supported by the Doris Duke Foundation (#2022092), the St. Jude Children’s Research Hospital Collaborative Research Consortium, the Harvard Stem Cell Institute, and the National Institutes of Health (UM1HG012010, R01HG013618, R01HL165061). L.P. was supported by a Rappaport MGH Research Scholar Award 2024-2029. S.L. holds a Banting Postdoctoral Fellowship from the Canadian Institutes of Health Research (CIHR). N.K. is supported by an Overseas Research Fellowship from the Japan Society for the Promotion of Science (JSPS). V.A.C.S. was supported by a Postdoctoral Fellowship from the American Heart Association (AHA) (25POST1377446). W.M. is supported by the National Institute of Health Research (NIH F30DK135340), and a Medical Scientist Training Fellowship from Harvard Stem Cell Institute. HSPCs were obtained from Fred Hutch Cooperative Center of Excellence in Hematology (U54DK106829). We thank Gabriele Casirati, Pietro Genovese, and Christian Brendel for helpful discussions, the HSCI-BCH Flow Cytometry Research Lab for technical support, and Bin Liu, Wen Xue, and Erik J. Sontheimer for communication of unpublished results.

## Author contributions

Conceptualization, S.L. and D.E.B; Methodology, S.L. and D.E.B; Investigation, S.L., N.K., G.H., B.B., V.A.C.S., W.M., L.H., L.P., S.A., D.E.B.; Original Draft, S.L.; Writing, Review, and Editing, S.L. and D.E.B.; Funding Acquisition, L.P., S.A., and D.E.B.; Supervision, L.P., S.A., and D.E.B.

---

**Supplementary Figure 1.**
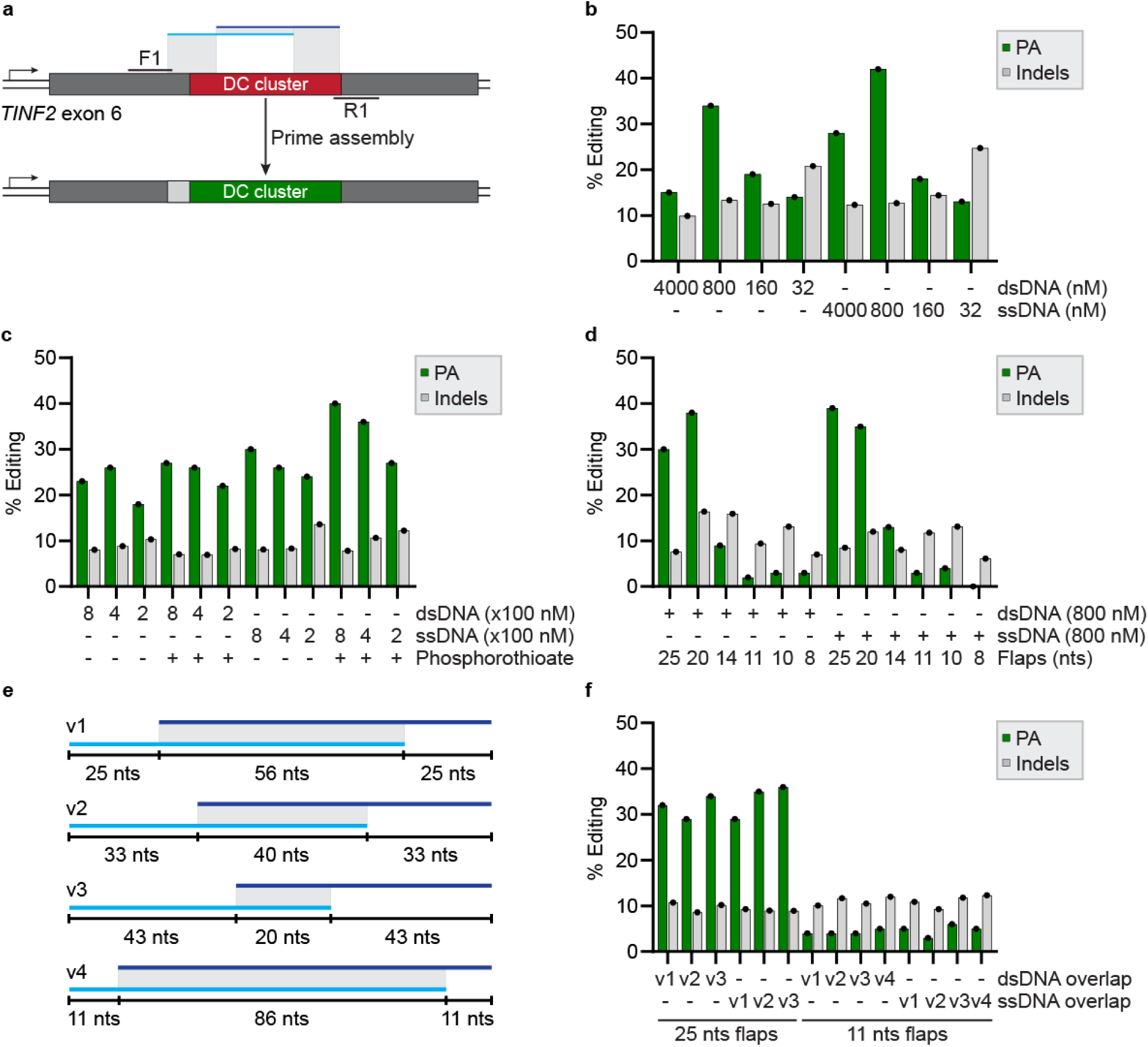
Prime assembly enables exon recoding at the *TINF2* dyskeratosis congenita cluster (Related to Fig. 1). **(a)** Schematic of *TINF2* DC cluster recoding using PA. (**b**) PA and indel quantification at *TINF2* as determined by ICE and TIDE analysis from Sanger sequencing, respectively. K562 cells were electroporated with PA vectors and the indicated concentration of donors with (dsDNA) or without (ssDNA) prior annealing, and genomic DNA was harvested 3 days post-nucleofection. *n* = 1 biological replicate. (**c**) Same as in (**b**), comparing donors with or without three phosphorothioate bonds at both the 5’ and 3’ ends. *n* = 1 biological replicate. (**d**) Same as in (**b**) using different flap lengths. *n* = 1 biological replicate. (**e**) Schematic representation of four different ssDNA designs with varying overlap lengths. (**f**) Same as in (**b**) using different ssDNA overlap designs shown in (**e**). *n* = 1 biological replicate.

**Supplementary Figure 2.**
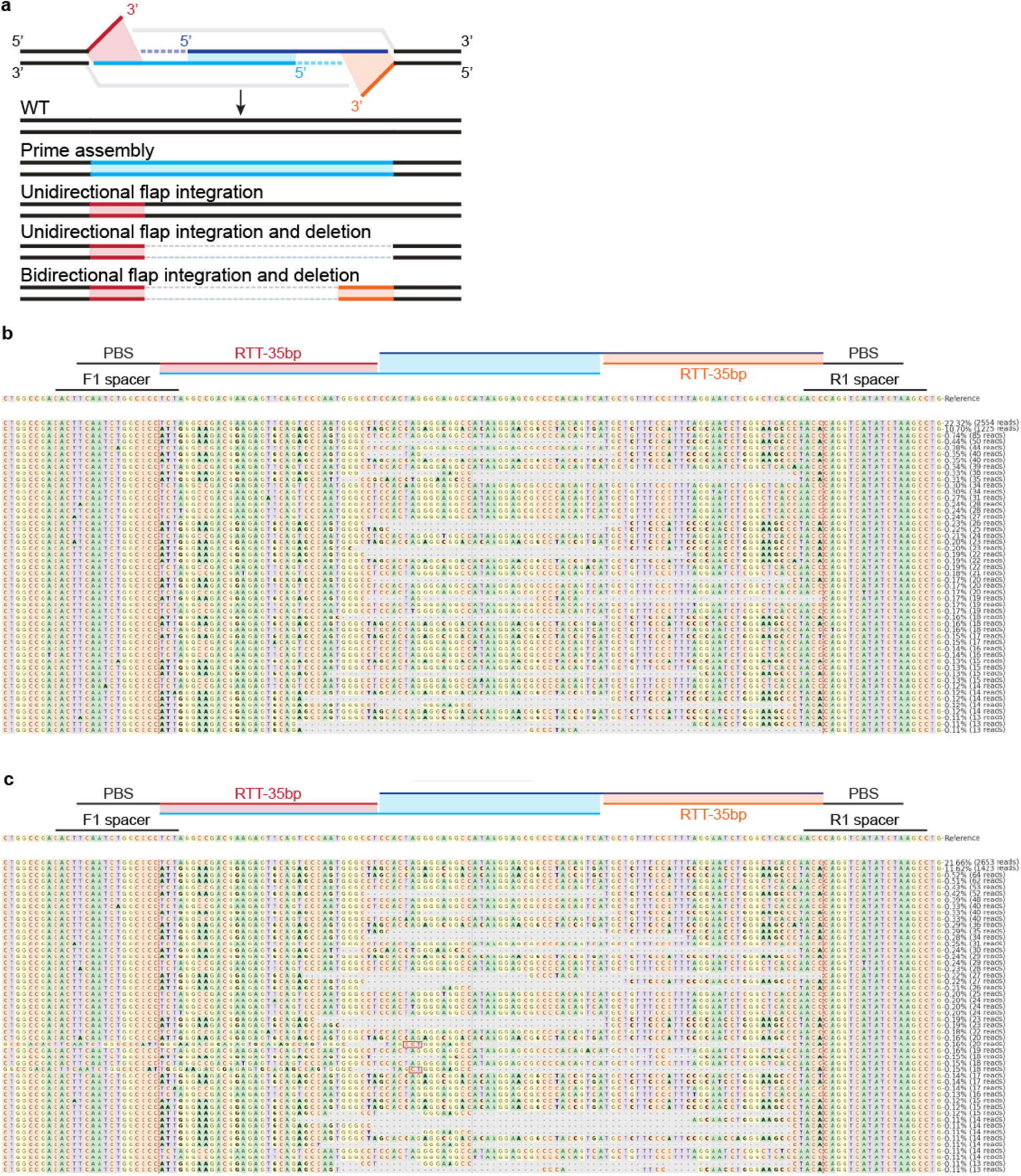
Characterization of prime assembly outcomes via amplicon sequencing (related to Fig. 1). (**a**) Schematic representation of the different types of outcomes observed after prime assembly as determined by amplicon sequencing. (**b**) Representative CRISPResso2 allele plot from Fig. 1c with 35 nucleotides flaps and 800 nM annealed (dsDNA) donor. Representative allele plot is from one out of three independent biological replicates. (**c**) Same as in (**b**) with 35 nucleotide flaps and 800 nM ssDNA donor. Representative allele plot is from one out of three independent biological replicates.

**Supplementary Figure 3.**
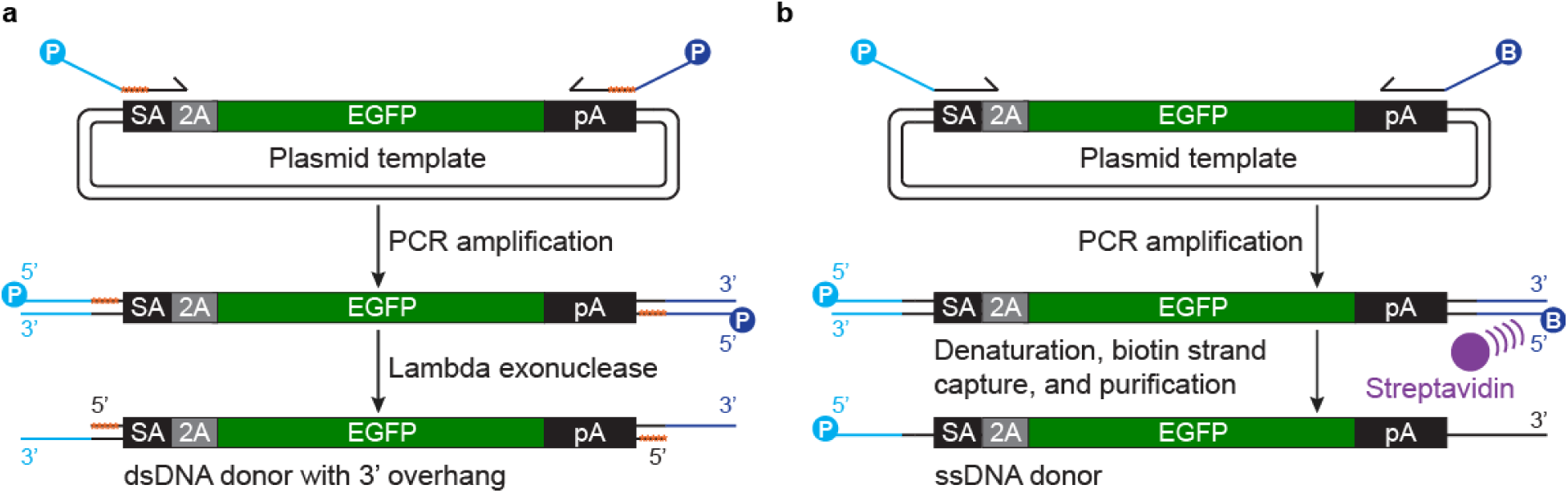
Generation of 3’-overhang dsDNA or ssDNA donors. (**a**) Schematic representation of the generation of 3’-overhang dsDNA donor. A plasmid of interest is used as a template to amplify the donor via PCR. The primers used for PCR include five phosphorothioate bonds (highlighted as orange stars) to protect the donor from lambda exonuclease digestion, 3’ overhangs of ≥ 20 nucleotides, and they are phosphorylated on the 5’ end. The dsDNA donor is then digested with lambda exonuclease to generate 3’ overhangs complementary to the prime assembly 3’ flaps. (**b**) Schematic representation of the generation of an ssDNA donor. A plasmid of interest is used as a template to amplify the donor via PCR. The primer used to amplify the ssDNA donor strand is phosphorylated on its 5’ end, and the primer used for the complementary ssDNA strand harbors a biotin on its 5’ end. After PCR amplification, the dsDNA PCR product is denatured and the biotinylated strand is immobilized on a magnet using streptavidin magnetic beads, and the ssDNA donor strand of interest is separated, neutralized, and purified.

**Supplementary Figure 4.**
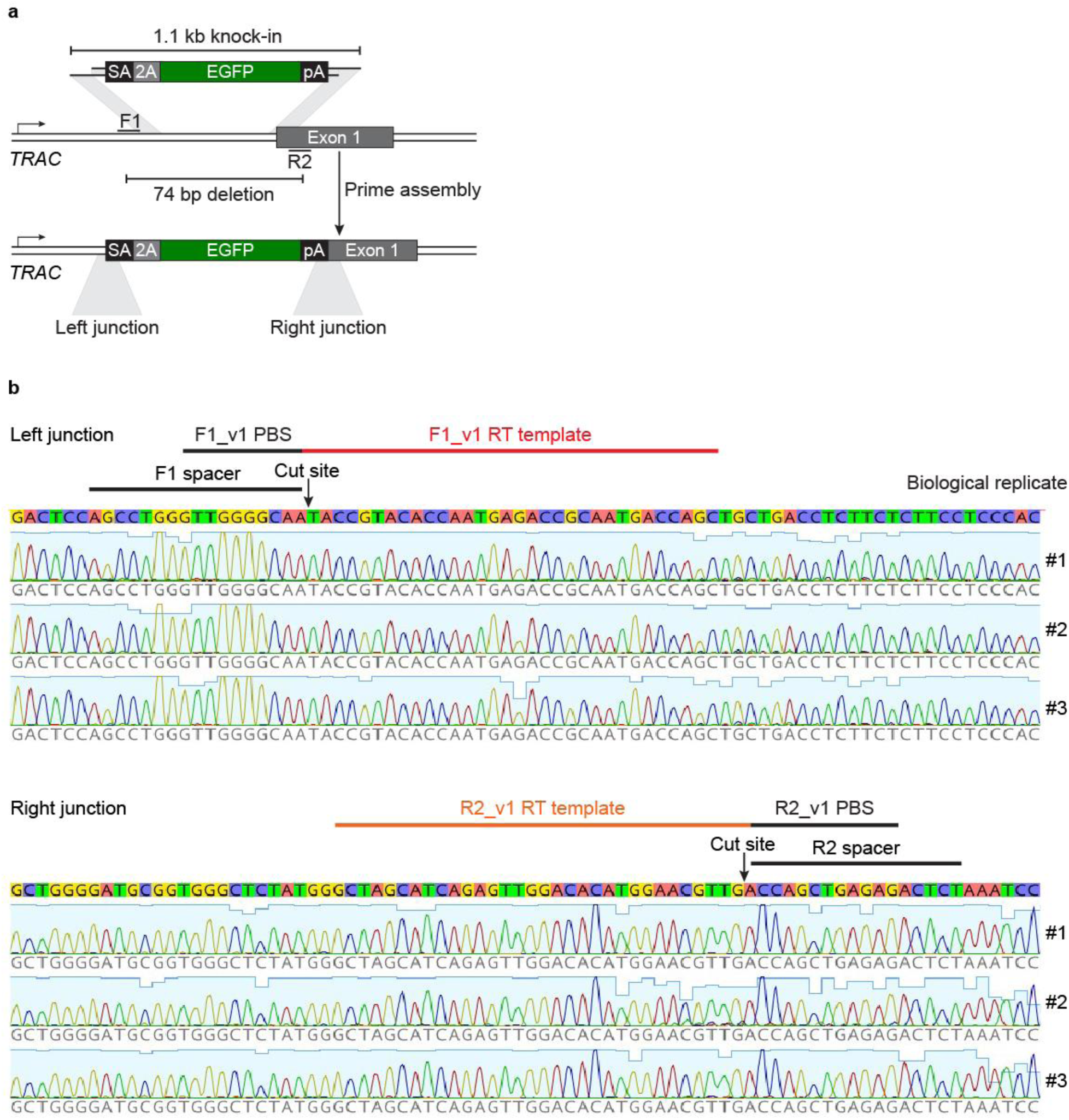
Prime assembly enables precise site-specific transgene integration (related to Fig. 2). (**a**) Schematic representation of targeted transgene integration at the *TRAC* locus using PA. (**b**) Representative Sanger chromatograms of the prime assembly junctions at the *TRAC* locus from the samples in Fig. 2b. *n* = 3 independent biological replicates.

**Supplementary Figure 5.**
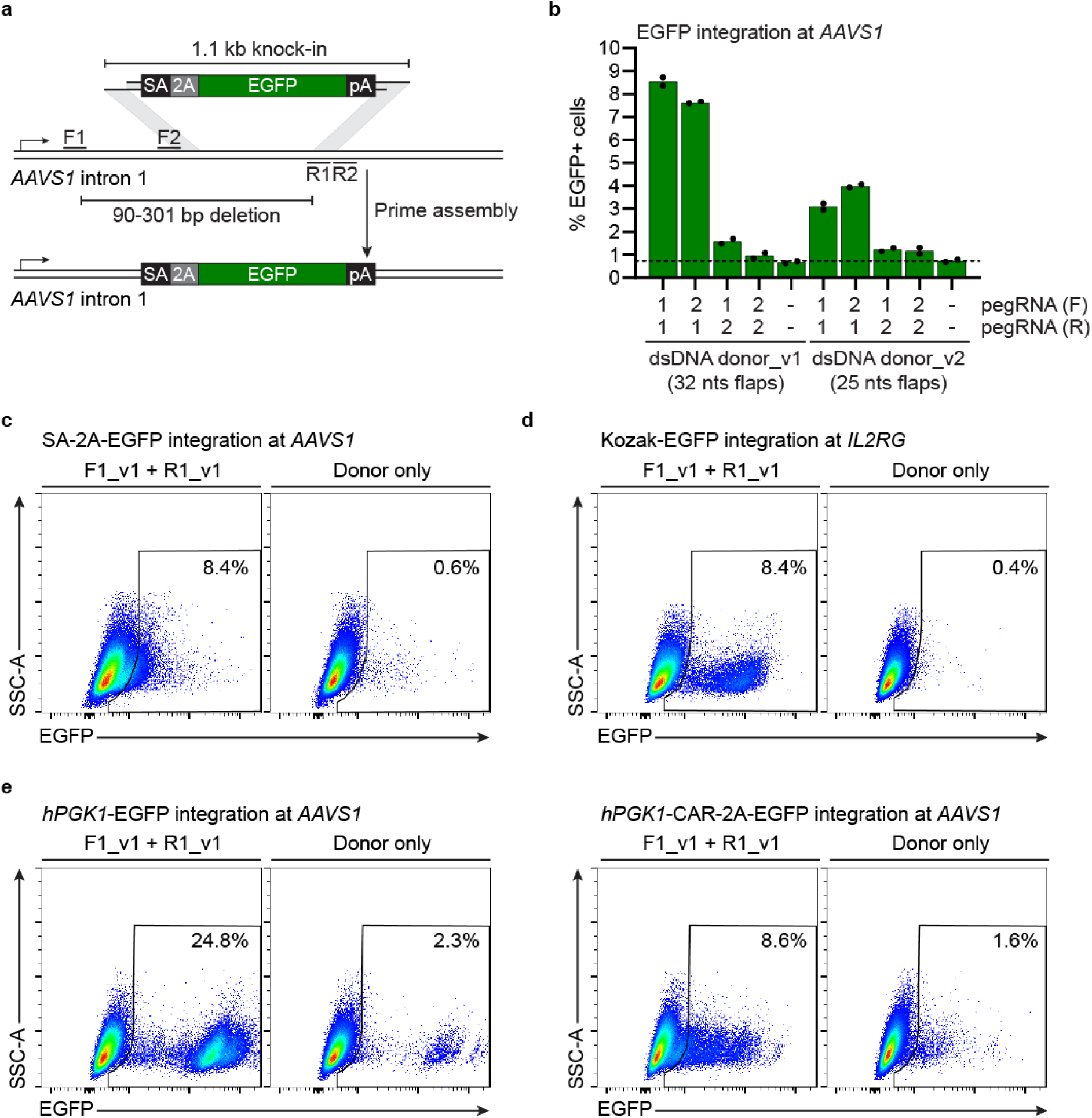
Functional site-specific reporter integration at *AAVS1* and *IL2RG* using 3’-overhang double-stranded DNA donors (related to Fig. 2). (**a**) Schematic representation of targeted transgene integration at the *AAVS1* locus using PA. (**b**) Percentage of EGFP^+^ cells as determined by flow cytometry. K562 cells were electroporated with PA vectors and 1.5 µg 3’-overhang dsDNA donor targeting *AAVS1*, and the percentage of EGFP^+^ cells was quantified 7 days post-nucleofection. *n* = 2 independent biological replicates. Dotted lines indicate the background level of the donor only controls. Dotted lines indicate the average background level of the donor only controls. (**c**) Representative FACS plots from the samples shown in (**b**). Representative FACS plots are from one out of two independent biological replicates. (**d**) Representative FACS plots from the samples shown in Fig. 2d. Representative FACS plots are from one out of three independent biological replicates. (**e**) Representative FACS plots from the samples shown in Fig. 2f. Representative FACS plots are from one out of three independent biological replicates.

**Supplementary Figure 6.**
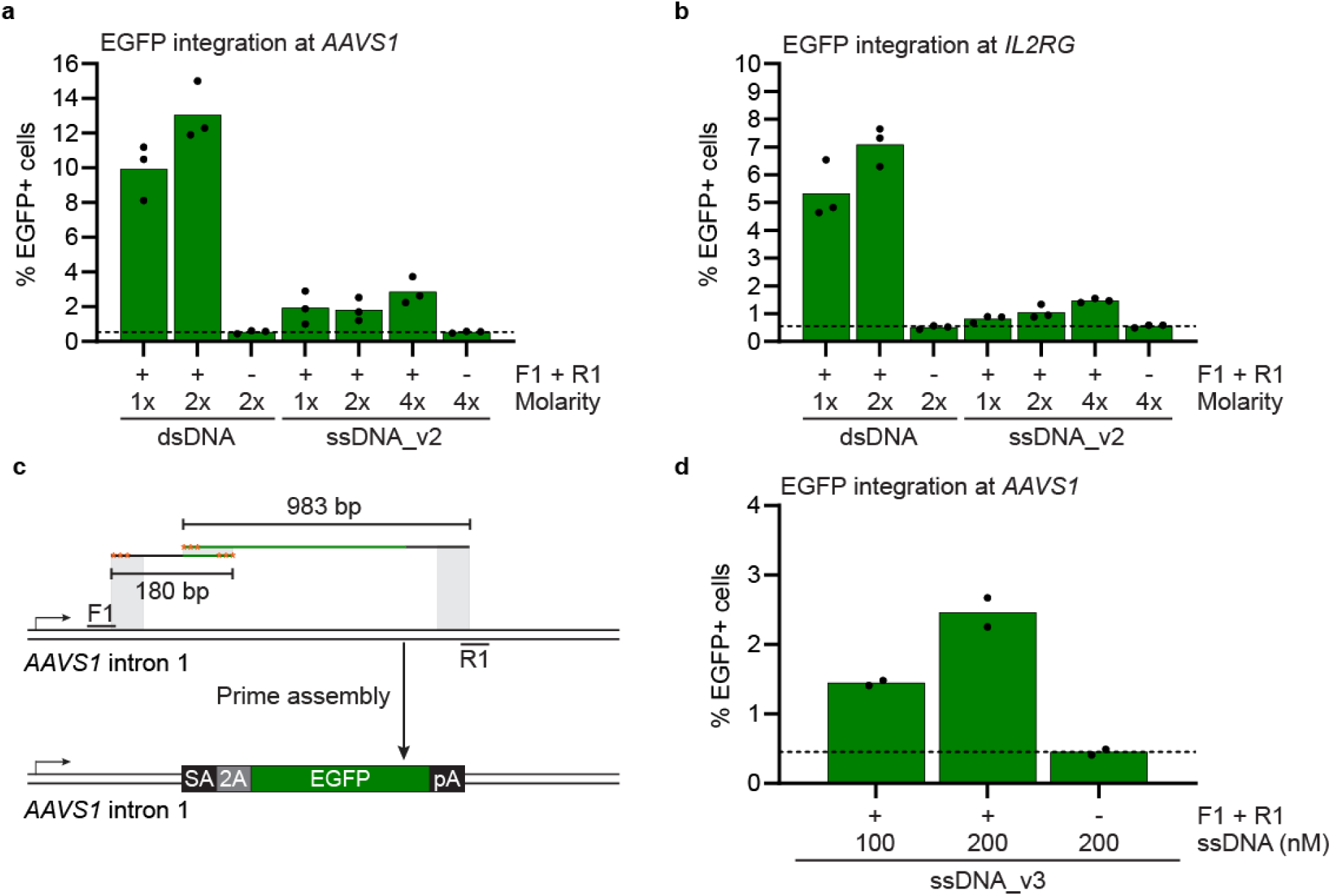
Site-specific transgene integration at *AAVS1* and *IL2RG* using increasing concentrations of single-stranded DNA donors (related to Figure 3). (**a**) Percentage of EGFP^+^ cells as determined by flow cytometry. K562 cells were electroporated with PA vectors and 1.5 µg 3’-overhang dsDNA donor (1X) targeting *AAVS1* or the indicated molar ratio of ssDNA donors, and the percentage of EGFP-expressing cells was quantified 7 days post-nucleofection. *n* = 3 independent biological replicates. Dotted lines indicate the average background level of the donor only controls. (**b**) Same as in (**a**) with donors targeting *IL2RG*. *n* = 3 independent biological replicates. (**c**) Schematic representation of targeted transgene integration at the *AAVS1* locus using ssDNA donors. Stabilizing phosphorothioate bonds are illustrated with orange stars. (**d**) Percentage of EGFP^+^ cells as determined by flow cytometry. K562 cells were electroporated with PA vectors and the indicated concentration of ssDNA donors targeting *AAVS1*, and the percentage of EGFP^+^ cells was quantified 7 days post-nucleofection. *n* = 2 independent biological replicates. Dotted lines indicate the average background level of the donor only controls.

**Supplementary Figure 7.**
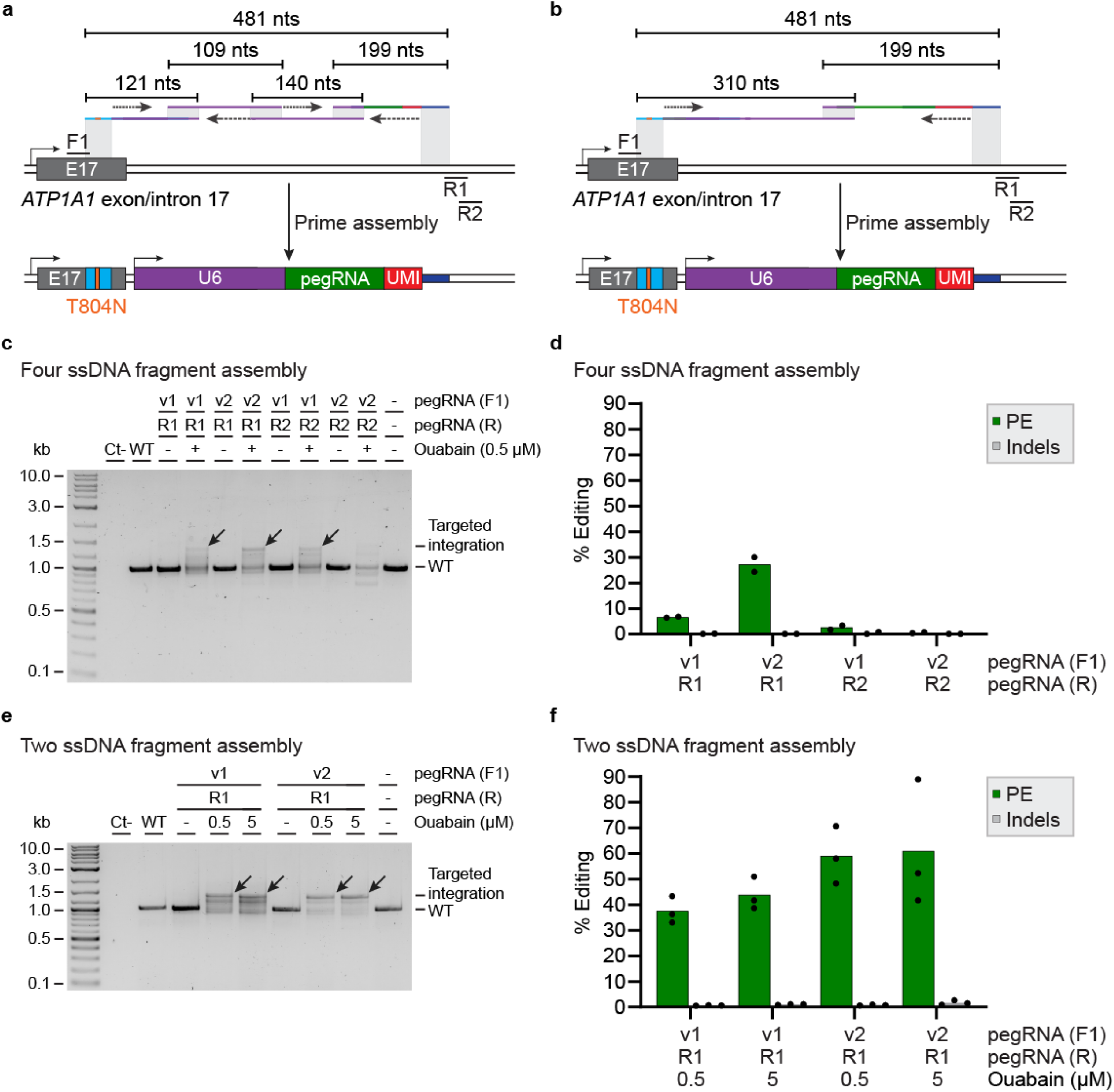
Targeted assembly and integration of up to four single-stranded DNA fragments in human cells. **(a)** Schematic representation of the installation of the ouabain resistance T804N mutation and the integration of a U6-pegRNA expression cassette at *ATP1A1* using four ssDNA donors. The arrows indicate fill-in synthesis. (**b**) Same as in (**a**) using two ssDNA donors. (**c**) PCR-based genotyping of site-specific *B2M*-targeting pegRNA expression cassette integration at *ATP1A1* using four ssDNA donors. K562 cells were electroporated with PA vectors and 800 nM of each ssDNA donor, and treated three days post-nucleofection with 0.5 µM ouabain until all non-resistant cells were eliminated. Representative gel is from one out of two independent biological replicates. (**d**) Prime editing and indel quantification at *B2M* as determined by BEAT and TIDE analysis from Sanger sequencing, respectively. Ouabain-resistant cells from (**c**) were re-electroporated with a prime editor vector and genomic DNA was harvested three days post-nucleofection. *n* = 2 independent biological replicates. (**e**) Same as in (**c**) for site-specific integration using two ssDNA donors. Representative gel is from one out of three independent biological replicates. (**f**) Same as in (**d**) with ouabain-resistant cells from (**e**). *n* = 3 independent biological replicates.

**Supplementary Figure 8.**
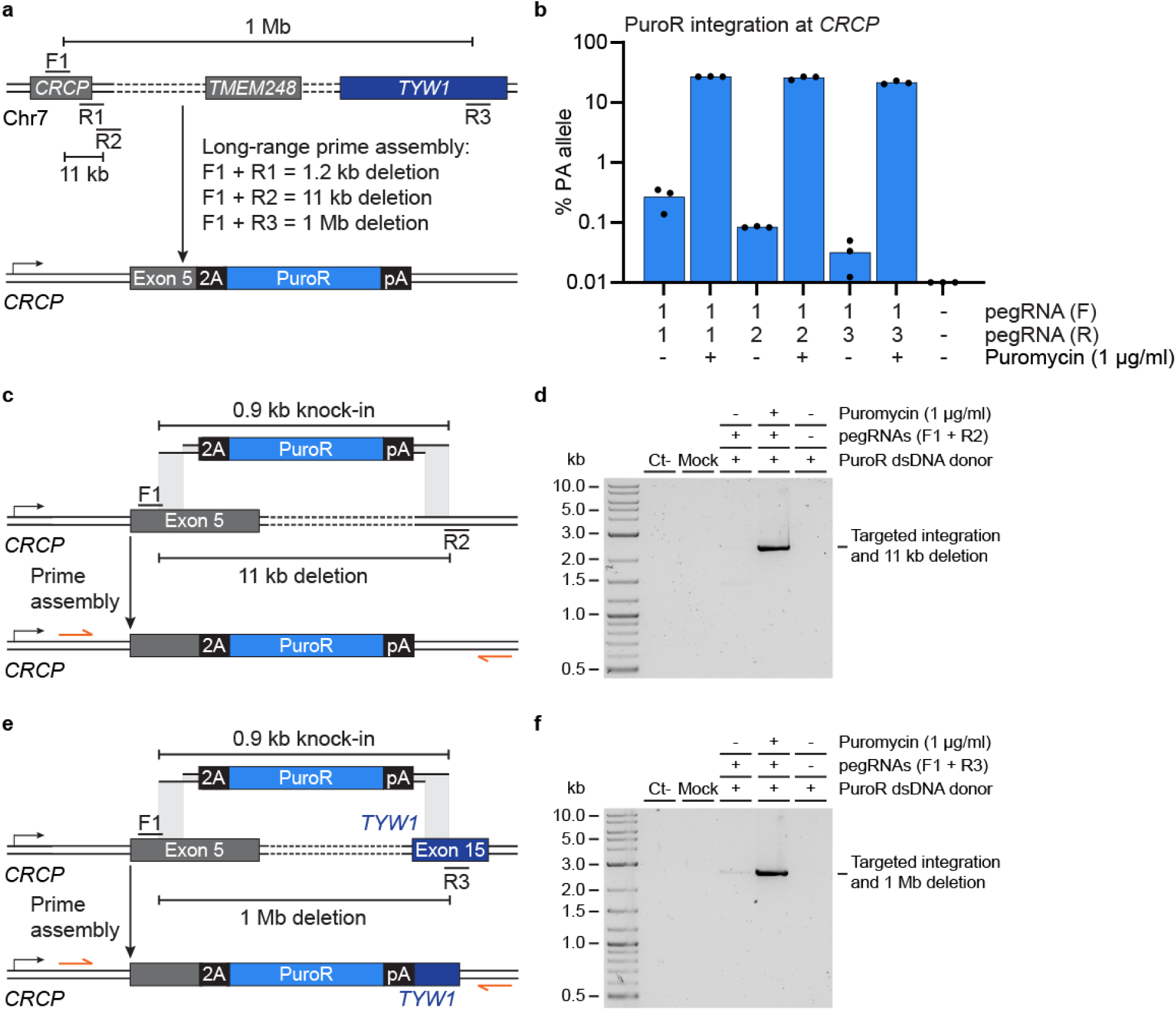
Long-range prime assembly enables megabase deletion at chromosome 7 in human cells (Related to Fig. 4). (**a**) Schematic representation of targeted megabase deletion at chromosome 7 and the integration of a puromycin-resistance (PuroR) cassette via long-range prime assembly. (**b**) Percentage of PA allele as determined by ddPCR. K562 cells were electroporated with PA vectors and 1.5 µg 3’ overhang dsDNA donor targeting *CRCP*, and cells were cultured with 1 µg/ml puromycin three days post-nucleofection until all non-resistant cells were eliminated. *n* = 3 independent biological replicates. (**c**) Schematic representation of targeted PuroR transgene integration and 11 kb deletion at the *CRCP* locus. (**d**) PCR-based genotyping of the expected PuroR transgene integration and 11 kb deletion allele after prime assembly and puromycin selection, as described in (**b**). Representative gel is from one out of three independent biological replicates. (**e**) Same as in (**c**) for targeted PuroR transgene integration and 1 Mb deletion at the *CRCP* locus. (**f**) Same as in (**d**) for PCR-based genotyping of the expected PuroR transgene integration and 1 Mb deletion allele after prime assembly and puromycin selection, as described in (**b**). Representative gel is from one out of three independent biological replicates.

**Supplementary Figure 9.**
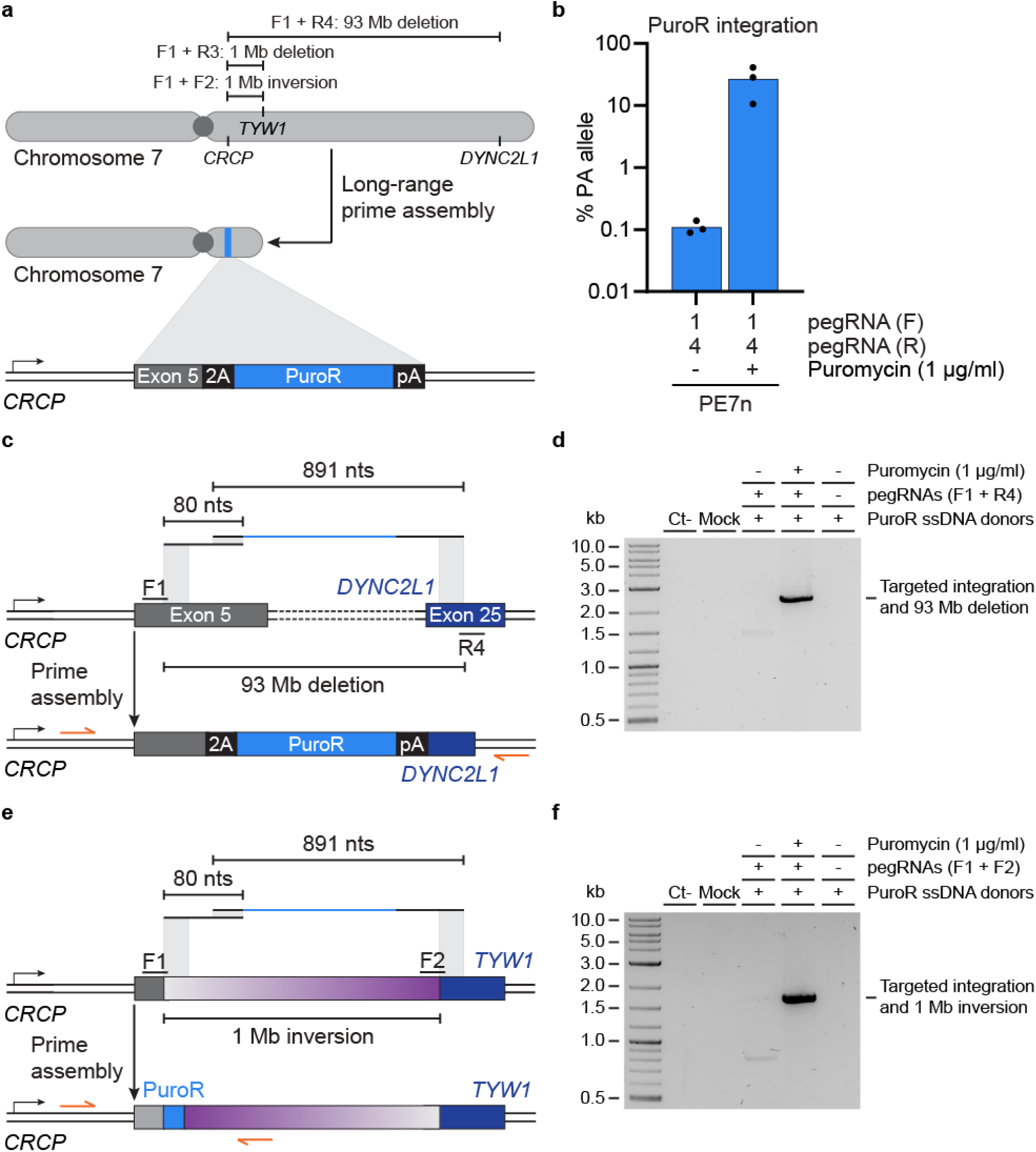
Long-range nuclease prime assembly enables targeted chromosome arm deletion and megabase inversion (related to figure 4). (**a**) Schematic representation of targeted PuroR transgene integration and the deletion of the q arm of chromosome 7. (**b**) Percentage of PA allele as determined by ddPCR. K562 cells were electroporated with nuclease PA vectors and 200 nM ssDNA donors targeting *CRCP*, and cells were cultured with 1 µg/ml puromycin three days post-nucleofection until all non-resistant cells were eliminated. *n* = 3 independent biological replicates. (**c**) Schematic representation of targeted PuroR transgene integration and the 93 Mb deletion at chromosome 7 using ssDNA donors. (**d**) PCR-based genotyping of the expected PuroR transgene integration and megabase deletion allele after prime assembly and puromycin selection, as described in (**b**). Representative gel is from one out of three independent biological replicates. (**e**) Schematic representation of targeted PuroR transgene integration and the 1 Mb inversion at chromosome 7 using ssDNA donors. (**f**) PCR-based genotyping of the expected PuroR transgene integration and megabase deletion allele after prime assembly and puromycin selection, as described in (**b**). Representative gel is from one experiment.

**Supplementary Figure 10.**
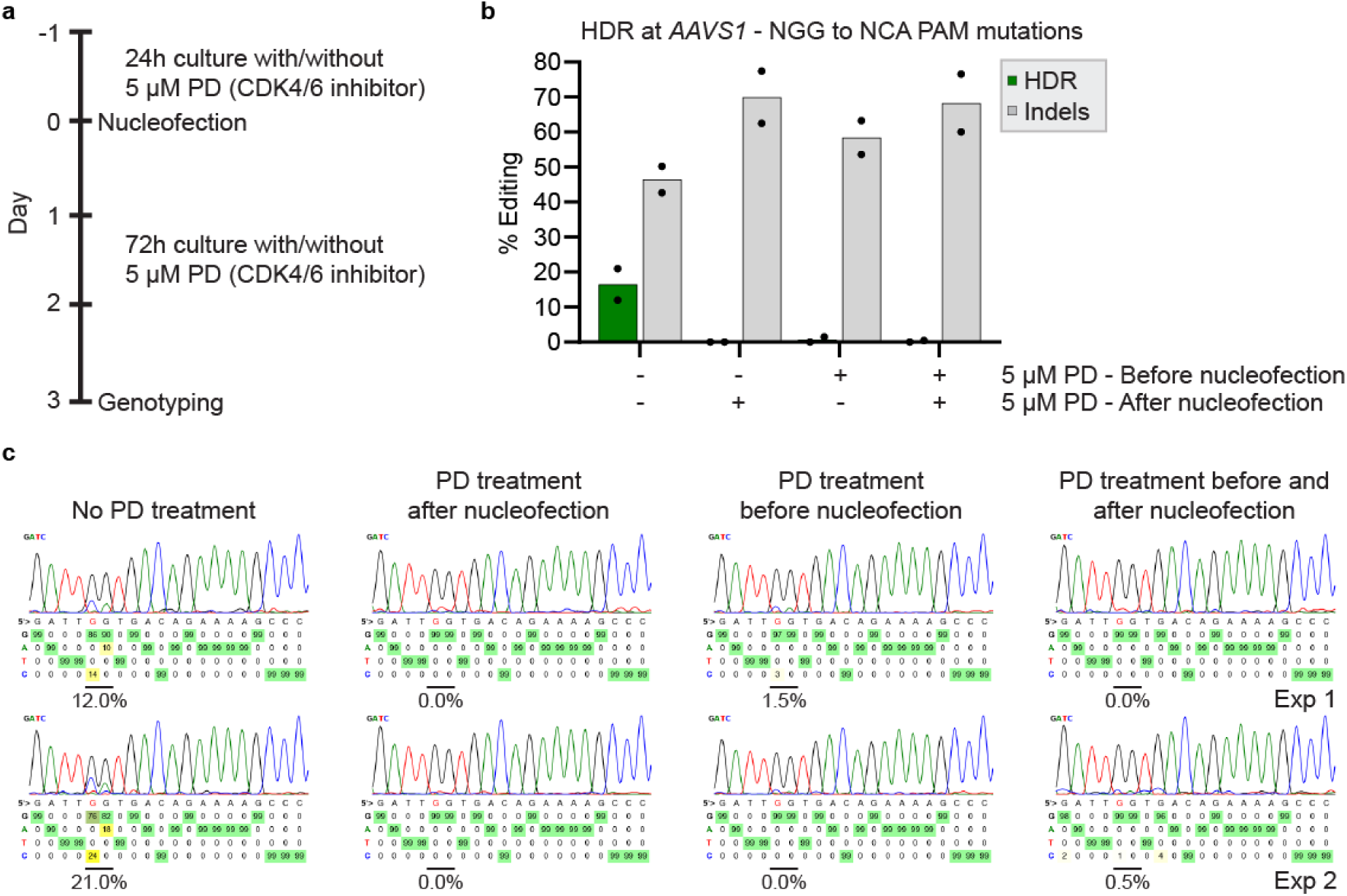
Inhibition of cell cycle progression abrogates HDR at the *AAVS1* locus in K562 cells (related to figure 5). (**a**) Timeline for the cell cycle experiment. K562 cells are cultured in the presence or absence of 5 µM PD 0332991, electroporated, and cultured for three days in the presence or absence of 5 µM PD 0332991. (**b**) HDR and indels quantification as determined by BEAT and TIDE analysis from Sanger sequencing, respectively. After 24 hours of culture in the presence or absence of 5 µM PD 0332991, K562 cells were electroporated with an SpCas9-sgRNA vector and an ssODN donor targeting *AAVS1*. Cells were cultured in the presence or absence of 5 µM PD 0332991 for 72 hours, and genomic DNA was harvested. *n* = 2 independent biological replicates. (**c**) Sanger (BEAT) allele plots from the two independent biological replicates shown in (**b**).

**Supplementary Figure 11.**
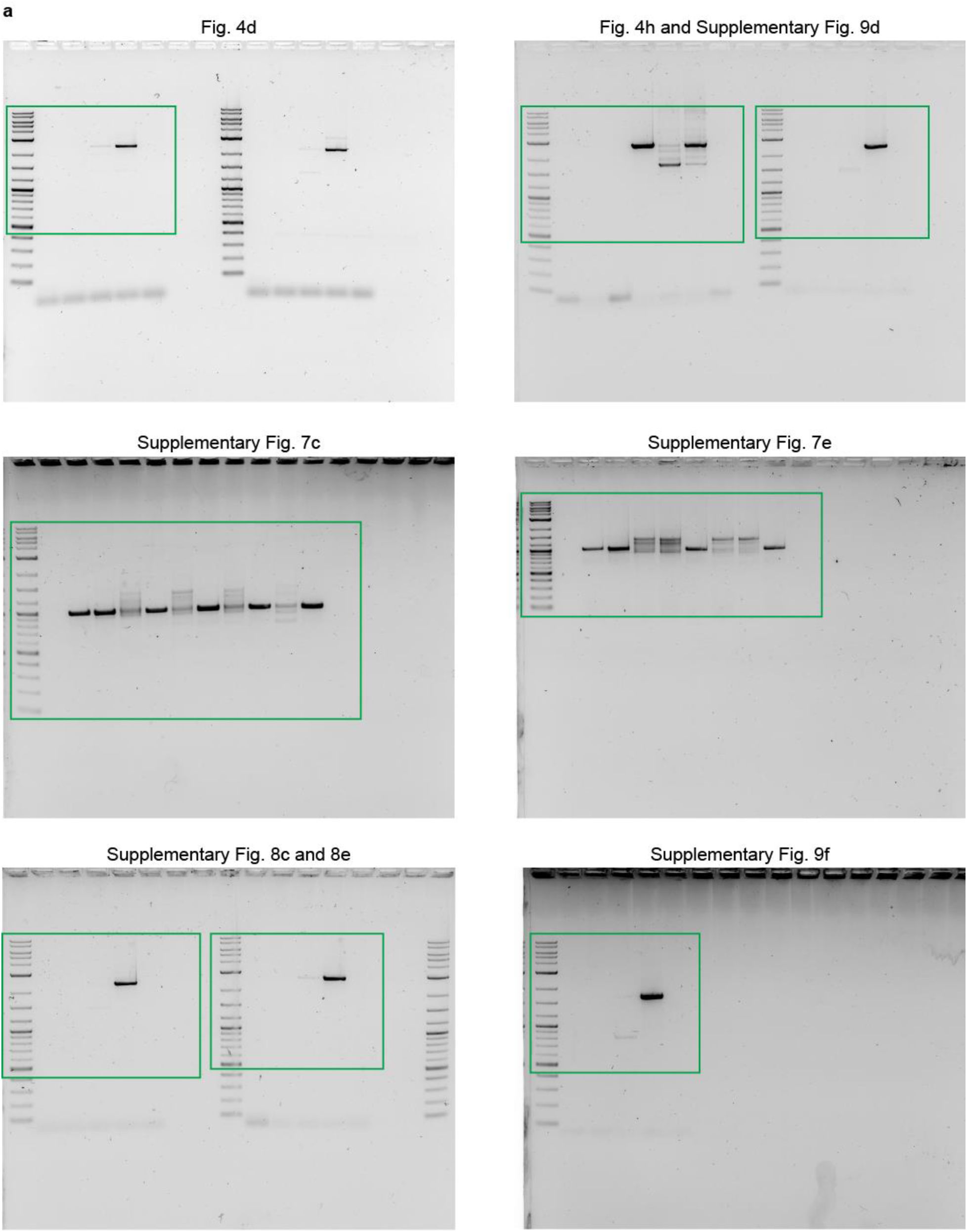
Uncropped scans of all gels from this study. (**a**) Cropping boundaries from each gel are highlighted in green.

## References

1. Tou, C. J. & Kleinstiver, B. P. Recent advances in double-strand break-free kilobase-scale genome editing technologies. Biochemistry 62, 3493–3499 (2023).

2. Eyquem, J. et al. Targeting a CAR to the TRAC locus with CRISPR/Cas9 enhances tumour rejection. Nature 543, 113–117 (2017).

3. Ellis, G. I., Sheppard, N. C. & Riley, J. L. Genetic engineering of T cells for immunotherapy. Nat Rev Genet 16, 103–107 (2021).

4. Mansilla-soto, J. et al. HLA-independent T cell receptors for targeting tumors with low antigen density. Nat Med 28, 345–352 (2022).

5. Mackensen, A. et al. Anti-CD19 CAR T cell therapy for refractory systemic lupus erythematosus. Nat Med 28, 2124–2132 (2022).

6. Bauer, D. E. et al. An erythroid enhancer of BCL11A subject to genetic variation determines fetal hemoglobin level. Science 342, 253–258 (2013).

7. Canver, M. C. et al. BCL11A enhancer dissection by Cas9-mediated in situ saturating mutagenesis. Nature 527, 192–197 (2015).

8. Wu, Y. et al. Highly efficient therapeutic gene editing of human hematopoietic stem cells. Nat Med 25, 776–783 (2019).

9. Frangoul, H. et al. Exagamglogene autotemcel for severe sickle cell disease. N Engl J Med 390, 1649–1662 (2024).

10. Locatelli, F. et al. Exagamglogene autotemcel for transfusion-dependent β-thalassemia. N Engl J Med 390, 1663–1676 (2024).

11. Hustedt, N. & Durocher, D. The control of DNA repair by the cell cycle. Nat Cell Biol 19, 1–9 (2017).

12. Murray, J. M. & Carr, A. M. Integrating DNA damage repair with the cell cycle. Curr Opin Cell Biol 52, 120–125 (2018).

13. Tsai, S. Q. et al. GUIDE-seq enables genome-wide profiling of off-target cleavage by CRISPR-Cas nucleases. Nat Biotechnol 33, 187–198 (2015).

14. Wienert, B. et al. Unbiased detection of CRISPR off-targets in vivo using DISCOVER-Seq. Science 364, 286–289 (2019).

15. Lazzarotto, C. R. et al. CHANGE-seq reveals genetic and epigenetic effects on CRISPR-Cas9 genome-wide activity. Nat Biotechnol 38, 1317–1327 (2020).

16. Cancellieri, S. et al. Human genetic diversity alters therapeutic gene editing off-target outcomes. Nat Genet 55, 34–43 (2023).

17. Kosicki, M. & Bradley, A. Repair of CRISPR–Cas9-induced double-stranded breaks leads to large deletions and complex rearrangements. Nat Biotechnol 36, 765–771 (2018).

18. Leibowitz, M. et al. Chromothripsis as an on-target consequence of CRISPR-Cas9 genome editing. Nat Genet 53, 895–905 (2021).

19. Nahmad, A. D. et al. Frequent aneuploidy in primary human T cells following CRISPR-Cas9 cleavage. Nat Biotechnol 40, 1807–1813 (2022).

20. Park, S. H. et al. Comprehensive analysis and accurate quantification of unintended large gene modifications induced by CRISPR-Cas9 gene editing. Sci Adv 8, eabo7676 (2022).

21. Tsuchida, C. A. et al. Mitigation of chromosome loss in clinical CRISPR-Cas9-engineered T cells. Cell 186, 4567–4582.e20 (2023).

22. Zeng, J. et al. Gene editing without ex vivo culture evades genotoxicity in human hematopoietic stem cells. Cell Stem Cell 32, 191–208 (2025).

23. Komor, A. C., Kim, Y. B., Packer, M. S., Zuris, J. A. & Liu, D. R. Programmable editing of a target base in genomic DNA without double-stranded DNA cleavage. Nature 533, 420–424 (2016).

24. Gaudelli, N. M. et al. Programmable base editing of A•T to G•C in genomic DNA without DNA cleavage. Nature 551, 464–471 (2017).

25. Anzalone, A. V. et al. Search-and-replace genome editing without double-strand breaks or donor DNA. Nature 576, 149–157 (2019).

26. Liu, B. et al. Targeted genome editing with a DNA-dependent DNA polymerase and exogenous DNA-containing templates. Nat Biotechnol 42, 1039–1045 (2024).

27. Silva, J. F. da et al. Click editing enables programmable genome writing using DNA polymerases and HUH endonucleases. Nat Biotechnol (2024) doi:10.1038/s41587-024-02324-x.

28. Kim, D. Y., Moon, S. Bin, Ko, J. H., Kim, Y. S. & Kim, D. Unbiased investigation of specificities of prime editing systems in human cells. Nucleic Acids Res 48, 10576–10589 (2020).

29. Liang, S., et al. Genome-wide profiling of prime editor off-target sites in vitro and in vivo using PE-tag. Nat Methods 20, 898–907 (2023).

30. Everette, K. A. et al. Ex vivo prime editing of patient haematopoietic stem cells rescues sickle-cell disease phenotypes after engraftment in mice. Nat Biomed Eng 7, 616–628 (2023).

31. Levesque, S. & Bauer, D. E. Gene correction for sickle cell disease hits its prime. Nat Biomed Eng 7, 605–606 (2023).

32. Fiumara, M. et al. Genotoxic effects of base and prime editing in human hematopoietic stem cells. Nat Biotechnol 42, 877–891 (2024).

33. Chen, P. J. & Liu, D. R. Prime editing for precise and highly versatile genome manipulation. Nat Rev Genet 24, 161–177 (2022).

34. Anzalone, A. V. et al. Programmable deletion, replacement, integration and inversion of large DNA sequences with twin prime editing. Nat Biotechnol 40, 731–740 (2022).

35. Yarnall, M. T. N. et al. Drag-and-drop genome insertion of large sequences without double-strand DNA cleavage using CRISPR-directed integrases. Nat Biotechnol 41, 500–512 (2022).

36. Pandey, S. et al. Efficient site-specific integration of large genes in mammalian cells via continuously evolved recombinases and prime editing. Nat Biomed Eng 9, 22–39 (2025).

37. Durrant, M. G. et al. Systematic discovery of recombinases for efficient integration of large DNA sequences into the human genome. Nat Biotechnol 41, 488–499 (2023).

38. Mukhametzyanova, L. et al. Activation of recombinases at specific DNA loci by zinc-finger domain insertions. Nat Biotechnol 42, 1844–1854 (2024).

39. Lampe, G. D. et al. Targeted DNA integration in human cells without double-strand breaks using CRISPR-associated transposases. Nat Biotechnol 42, 87–98 (2024).

40. Witte, I. P. et al. Programmable gene insertion in human cells with a laboratory-evolved CRISPR-associated transposase. Science 388, eadt5199 (2025).

41. Perry, N. T., et al. Megabase-scale human genome rearrangement with programmable bridge recombinases. BioRxiv. Preprint at https://www.biorxiv.org/content/10.110 (2025).

42. Estes, B. J. G. et al. Development of circular AAV cargos for targeted seamless insertion with large serine integrases. Mol Ther Methods Clin Dev 33, 101490 (2025).

43. Charlesworth, C. T., Hsu, I., Wilkinson, A. C. & Nakauchi, H. Immunological barriers to haematopoietic stem cell gene therapy. Nat Rev Immunol 22, 719–733 (2022).

44. Decout, A., Katz, J. D., Venkatraman, S. & Ablasser, A. The cGAS–STING pathway as a therapeutic target in inflammatory diseases. Nat Rev Immunol 21, 548–569 (2021).

45. Ferrari, S. et al. Choice of template delivery mitigates the genotoxic risk and adverse impact of editing in human hematopoietic stem cells. Cell Stem Cell 29, 1428–1444.e9 (2022).

46. Bauer, D. E. Pervasive donor DNA integration defies precision gene editing of hematopoietic stem cells. Cell Stem Cell 29, 1426–1427 (2022).

47. Schiroli, G. et al. Precise gene editing preserves hematopoietic stem cell function following transient p53-mediated DNA damage response. Cell Stem Cell 24, 551–565 (2019).

48. DeWitt, M. A. et al. Selection-free genome editing of the sickle mutation in human adult hematopoietic stem/progenitor cells. Sci Transl Med 8, 360ra134 (2016).

49. Shy, B. R. et al. High-yield genome engineering in primary cells using a hybrid ssDNA repair template and small-molecule cocktails. Nat Biotechnol 41, 521–531 (2023).

50. Xie, K. et al. Efficient non-viral immune cell engineering using circular single-stranded DNA-mediated genomic integration. Nat Biotechnol (2024) doi:10.1038/s41587-024-02504-9.

51. Gibson, D. G. et al. Enzymatic assembly of DNA molecules up to several hundred kilobases. Nat Methods 6, 343–345 (2009).

52. Jones, D. H. & Howard, B. H. A rapid method for recombination and site-specific mutagenesis by placing homologous ends on DNA using polymerase chain reaction. Biotechniques 10, 62–66 (1991).

53. Bubeck, P., Winkler, M. & Bautsch, W. Rapid cloning by homologous recombination in vivo. Nucleic Acids Res 21, 3601–3602 (1993).

54. Watson, J. F. & García-Nafría, J. In vivo DNA assembly using common laboratory bacteria: A re-emerging tool to simplify molecular cloning. Journal of Biological Chemistry 294, 15271–15281 (2019).

55. Kostylev, M., Otwell, A. E., Richardson, R. E. & Suzuki, Y. Cloning should be simple: Escherichia coli DH5α-mediated assembly of multiple DNA fragments with short end homologies. PLoS One 10, e0137466 (2015).

56. Huang, F., Spangler, J. R. & Huang, A. Y. In vivo cloning of up to 16 kb plasmids in E. coli is as simple as PCR. PLoS One 12, e0183974 (2017).

57. García-Nafría, J., Watson, J. F. & Greger, I. H. IVA cloning: A single-tube universal cloning system exploiting bacterial In Vivo Assembly. Sci Rep 6, 27459 (2016).

58. Wang, J. et al. Efficient targeted insertion of large DNA fragments without DNA donors. Nat Methods 19, 331–340 (2022).

59. Jiang, T., Zhang, X., Weng, Z. & Xue, W. Deletion and replacement of long genomic sequences using prime editing. Nat Biotechnol 40, 227–234 (2021).

60. Choi, J. et al. Precise genomic deletions using paired prime editing. Nat Biotechnol 40, 218–226 (2022).

61. Dokal, I., Vulliamy, T., Mason, P. & Bessler, M. Clinical utility gene card for: Dyskeratosis congenita-update 2015. European Journal of Human Genetics 23, e2–e4 (2015).

62. Choo, S. et al. Editing TINF2 as a potential therapeutic approach to restore telomere length in dyskeratosis congenita. Blood 140, 608–618 (2022).

63. Savage, S. A. et al. TINF2, a component of the shelterin telomere protection complex, is mutated in dyskeratosis congenita. Am J Hum Genet 82, 501–509 (2008).

64. Walne, A. J., Vulliamy, T., Beswick, R., Kirwan, M. & Dokal, I. TINF2 mutations result in very short telomeres: Analysis of a large cohort of patients with dyskeratosis congenita and related bone marrow failure syndromes. Blood 112, 3594–3600 (2008).

65. Alter, B. P. et al. Telomere length is associated with disease severity and declines with age in dyskeratosis congenita. Haematologica 97, 353–359 (2012).

66. Norris, K. et al. High-throughput STELA provides a rapid test for the diagnosis of telomere biology disorders. Hum Genet 140, 945–955 (2021).

67. Chen, P. J. et al. Enhanced prime editing systems by manipulating cellular determinants of editing outcomes. Cell 184, 5635–5652 (2021).

68. Yan, J. et al. Improving prime editing with an endogenous small RNA-binding protein. Nature 628, 639–647 (2024).

69. Anzalone, A. V. et al. Search-and-replace genome editing without double-strand breaks or donor DNA. Nature 576, 149–157 (2019).

70. Levesque, S. et al. Marker-free co-selection for successive rounds of prime editing in human cells. Nat Commun 13, 5909 (2022).

71. Yan, X. Efficient precise integration of large DNA sequences with 3’-overhang dsDNA donors using CRISPR/Cas9. Proceedings of the National Academy of Sciences 120, e2221127120 (2023).

72. Roth, T. L. et al. Reprogramming human T cell function and specificity with non-viral genome targeting. Nature 559, 405–409 (2018).

73. Levesque, S., Cosentino, A., Verma, A., Genovese, P. & Bauer, D. E. Enhancing prime editing in hematopoietic stem and progenitor cells by modulating nucleotide metabolism. Nat Biotechnol 43, 534–538 (2025).

74. Wright, B. W., Molloy, M. P. & Jaschke, P. R. Overlapping genes in natural and engineered genomes. Nat Rev Genet 23, 154–168 (2021).

75. Zikry, T. M. et al. Cell cycle plasticity underlies fractional resistance to palbociclib in ER+/HER2− breast tumor cells. Proceedings of the National Academy of Sciences 121, e2309261121 (2024).

76. Paquet, D. et al. Efficient introduction of specific homozygous and heterozygous mutations using CRISPR/Cas9. Nature 533, 125–129 (2016).

77. Kotini, A. G. & Papapetrou, E. P. Engineering of targeted megabase-scale deletions in human induced pluripotent stem cells. Exp Hematol 87, 25–32 (2020).

78. Essletzbichler, P. et al. Megabase-scale deletion using CRISPR/Cas9 to generate a fully haploid human cell line. Genome Res 24, 2059–2065 (2025).

79. Girish, V. et al. Oncogene-like addiction to aneuploidy in human cancers. Science 381, eadg4521 (2023).

80. Shin, J. J. et al. Controlled cycling and quiescence enables efficient HDR in engraftment-enriched adult hematopoietic stem and progenitor cells. Cell Rep 32, 108093 (2020).

81. Lauridsen, F. K. B. et al. Differences in cell cycle status underlie transcriptional heterogeneity in the HSC compartment. Cell Rep 24, 766–780 (2018).

82. Oedekoven, C. A. et al. Hematopoietic stem cells retain functional potential and molecular identity in hibernation cultures. Stem Cell Reports 16, 1614–1628 (2021).

83. Ferrari, S. et al. Efficient gene editing of human long-term hematopoietic stem cells validated by clonal tracking. Nat Biotechnol 38, 1298–1308 (2020).

84. Hanlon, K. S. et al. High levels of AAV vector integration into CRISPR-induced DNA breaks. Nat Commun 10, 4439 (2019).

85. Suchy, F. P. et al. Genome engineering with Cas9 and AAV repair templates generates frequent concatemeric insertions of viral vectors. Nat Biotechnol 43, 204–213 (2024).

86. Giuffrida, L. et al. CRISPR/Cas9 mediated deletion of the adenosine A2A receptor enhances CAR T cell efficacy. Nat Commun 12, 3236 (2021).

87. Carnevale, J. et al. RASA2 ablation in T cells boosts antigen sensitivity and long-term function. Nature 609, 174–182 (2022).

88. Wei, J. et al. Targeting REGNASE-1 programs long-lived effector T cells for cancer therapy. Nature 576, 471–476 (2019).

89. Nelson, J. W. et al. Engineered pegRNAs improve prime editing efficiency. Nat Biotechnol 40, 402–410 (2021).

90. Chen, B. et al. Dynamic imaging of genomic loci in living human cells by an optimized CRISPR/Cas system. Cell 155, 1479–1491 (2013).

91. Cong, L. et al. Multiplex genome engineering using CRISPR/Cas systems. Science 339, 819–823 (2013).

92. Casirati, G. et al. Epitope editing enables targeted immunotherapies for acute myeloid leukemia. Nature 621, 404–414 (2023).

93. Brinkman, E. K., Chen, T., Amendola, M. & Van Steensel, B. Easy quantitative assessment of genome editing by sequence trace decomposition. Nucleic Acids Res 42, e168 (2014).

94. Xu, L., Liu, Y. & Han, R. BEAT: A Python program to quantify base editing from Sanger sequencing. CRISPR J 2, 223–229 (2019).

95. Conant, D. et al. Inference of CRISPR edits from Sanger trace data. CRISPR J 5, 123–130 (2022).

96. Clement, K. et al. CRISPResso2 provides accurate and rapid genome editing sequence analysis. Nat Biotechnol 37, 215–226 (2019).

